# An automated pipeline for reconstructing whole genome duplications

**DOI:** 10.64898/2026.08.10.743943

**Authors:** Łukasz Niezabitowski, Anthony K. Redmond, Aoife McLysaght

## Abstract

Whole genome duplications leave lasting traces in our genomes. How these present in terms of gene content and order varies over time. While collinear blocks of paralogs, long stretches of conserved gene order and content termed ‘microsynteny’, are a distinctive feature of comparatively recent WGD and have been integral in reconstructing the history of ancestral duplication events, this signal degrades over time, making analysis of older events non-trivial. While gene order degrades quickly, gene content is often better conserved and recent work takes advantage of this to reconstruct older events and ancestral pre-WGD and post-WGD chromosomes. However, these new methods are complicated and not well-documented. Here we develop an automated and user-friendly pipeline for reconstructing ancestral chromosomes before and after WGD, and use the conservation of gene content to infer chromosomal rearrangement events in this timeframe. We verify the efficacy of our tool by reconstructing the ancestral acipenseriform, a model system for vertebrate WGD and rediploidisation. Our pipeline should serve to make ancestral reconstruction more accessible and provide a solid foundation for future analysis.

## Introduction

Whole genome duplications (WGDs) have occurred throughout the eukaryotic tree of life, with prominent examples in plants (The Arabidopsis Genome Initiative, 2000; Mandáková and Lysak, 2018; Clark and Donoghue, 2018), yeast (Wolfe and Shields, 1997), and vertebrates (McLysaght et al., 2002; Volff, 2005; Dehal and Boore, 2005; Nakatani et al., 2021). These events have been implicated in unlocking the evolutionary potential of lineages by providing raw genetic material and facilitating innovations and lineage diversity (Wolfe, 2001; Conant and Wolfe, 2008; Van de Peer et al., 2009, 2017; Mandáková and Lysak, 2018). Despite the variety in environmental conditions and evolutionary pressures surrounding WGD (Van de Peer et al., 2017), their impact on modern genomes persists for millions of years and can often be observed in the form of conserved gene order and content between formerly duplicated genomic regions (Sacerdot et al., 2018; Singh and Isambert, 2019; Simakov et al., 2020; Nakatani et al., 2021); Initially, gene order and content on duplicate regions will be near-identical. However, with evolutionary time, this pattern becomes degraded by chromosome fusion, fission, translocation, and loss events, as well as internal chromosomal rearrangements and gene loss events (Berthelot et al., 2014; Lien et al., 2016; Robertson et al., 2017; Simakov et al., 2020; Nakatani et al., 2021; Simakov et al., 2022; Redmond et al., 2023). Despite this, we can still uncover traces of the formerly symmetrically duplicated genome in the form of collinear blocks of paralogs (Wolfe and Shields, 1997; McLysaght et al., 2002; Kellis et al., 2004; Dehal and Boore, 2005; Sacerdot et al., 2018; Singh and Isambert, 2019; Simakov et al., 2020; Nakatani et al., 2021). This is true both on a ‘micro’ scale, describing the conservation of gene order and content in small genomic regions, as well as a ‘macro’ scale where gene order is broken due to extensive shuffling, but homologous segments remain enriched in paralogs (Sacerdot et al., 2018; Nakatani et al., 2021).

The significance and potential of these collinear blocks was quickly recognised (Ohno, 1970) and leveraged to better understand the WGDs that created them. Indeed, much of the evidence for well-studied WGD events in yeast, plants, and vertebrates, has originated from identifying and mapping the locations of these genomic regions (Wolfe and Shields, 1997; The Arabidopsis Genome Initiative, 2000; McLysaght et al., 2002; Dehal and Boore, 2005). Similarly, recent work has employed a microsynteny-based approach to build a comprehensive dataset of WGD duplicates. Rather than focusing on sequence similarity or phylogenetics (Dehal and Boore, 2005; Nakatani et al., 2007; Robertson et al., 2017), which were often used to identify ohnologs *ad hoc*, Singh et al. (2015) and Sacerdot et al. (2018) leverage the preservation and extensive coverage of homologous segments in many vertebrate genomes (Larkin et al., 2009; Venkatesh et al., 2014; Braasch et al., 2016; Lu et al., 2023). Of course, these advances have been possible, in no small part, thanks to the standardised tools and methods which allow researchers to quickly and easily identify these paralogous blocks, as well as an increasingly dense availability of high-quality genomes (Wang et al., 2012; Proost et al., 2012; Wang et al., 2024).

Presently, most of these tools focus on microsynteny. While this may be sufficient for solving some paleopolyploids, as well as reconsructing gene order in recently-diverged lineages (Muffato et al., 2023; Bernard et al., 2025), microsynteny degrades over time (Nakatani and McLysaght, 2017; Nakatani et al., 2021), making it less effective for distant comparisons or in genomes with high rates of chromosomal rearrangement, as is the case in many invertebrate genomes (Simakov et al., 2020; Nakatani et al., 2021). As such, recent advances in genome sequencing and the broad availability of chromosome-level assemblies have prompted the development of alternative methods, making better use of macrosynteny—the chromosome-wide conservation of homolog content (Schultz et al., 2023; Lewin et al., 2025). Recent work (Nakatani and McLysaght, 2017; Simakov et al., 2020; Nakatani et al., 2021; Niezabitowski et al., forthcoming) has taken advantage of this approach to reconstruct the ancestral complement of chromosomes before and after the vertebrate 2R, providing valuable insight into early vertebrate evolution and disentangling the previously contested history of 2R in vertebrates (Fried et al., 2003; Furlong et al., 2007; Mehta et al., 2013; Smith et al., 2013, 2018). Similar methods have also been applied to other WGD events, such as the teleost Ts3R (Nakatani and McLysaght, 2017; Parey et al., 2022), as well as for tracking the evolution of genomic segments in unduplicated genomes (Schultz et al., 2023), or reconstructing ancient animal chromosomal linkage groups (Simakov et al., 2022).

Unfortunately, unlike previous microsynteny-based approaches, these new tools are far from standardised or user-friendly. Some of these methods rely on the prior availability of high-quality ohnolog datasets or genomic resources. While this is not an obstacle for well-studied clades, these resources may not always be available, for example in non-marine annelids, where several rounds of putative WGD have recently been proposed (Lewin et al., 2024; Vargas-Chávez et al., 2025). Other approaches are not easy to use and not well-documented (Nakatani and McLysaght, 2017; Nakatani et al., 2021; Simakov et al., 2020). These tools often use *ad hoc* scripts to perform parts of the analysis, many of which are not publicly available, making the process laborious and time intensive. Importantly, many of these tools may not fully account for the impact of WGD on the evolutionary trajectories of homologous genomic regions. Given the unique genomic structure that arises after WGD and the resulting evolutionary opportunities, it is essential to identify and consider genomic segments originating by WGD.

Here we present an automated pipeline for reconstructing ancestral whole genome duplication events. We show that it is fast, accurate, and easily adaptable to a wide range of datasets. We reconstruct the ancestral acipenseriform genome, a major vertebrate lineage with an ancient WGD and a key model in recent work on asynchronous rediploidisation, to showcase our tool and verify its effectiveness. The pipeline is easy to install and easy to use, and was designed with the intention of allowing a broad range of researchers, from novices to expert users, to unlock the full potential of synteny analysis and paleopolyploidy insights. Much like the microsynteny-based tools and methods that enabled some of the early discoveries in WGD (McLysaght et al., 2002; Dehal and Boore, 2005; Proost et al., 2012; Singh and Isambert, 2019), this next generation of tools has the potential to resolve many difficult problems in evolutionary genomics.

## Results

### An overview of the WGD reconstruction pipeline and suggested workflow

Our pipeline aims to automate and standardise the workflow of reconstructing ancestral whole genome duplications in an open and readily useable way. The workflow, as illustrated in Fig. 1, involves: (i) generating orthologs and paralogs (Buchfink et al., 2021), (ii) segmenting modern genomes based on macrosynteny (Nakatani and McLysaght, 2017; Nakatani et al., 2021), (iii) reconstructing pre-WGD and post-WGD ancestral chromosomes, and (iv) inferring the chromosomal fusion and fission events that occurred around this time-frame (Nakatani et al., 2021). The required input (Fig. 1a) can be provided as genomic FASTA and general feature format (GFF) files. All other downstream files are automatically generated by the pipeline, but alternatively, can be provided by the researcher. This may be desirable in the case of proteomes (Fig. 1b) if genomic FASTAs or GFF files are not easily accessible, or in the case of homologs (Fig. 1c) if high-quality ortholog and paralog datasets already exist elsewhere. Pipeline outputs and intermediate files use well established bioinformatics file-formats (Pearson and Lipman, 1988; Eilbeck et al., 2005) and TSV files in the case of tabulated data. This maximises inter-operability with other bioinformatics tools. These data can be visualised using scripts included with the pipeline and used in further downstream analyses.

**Fig. 1.**
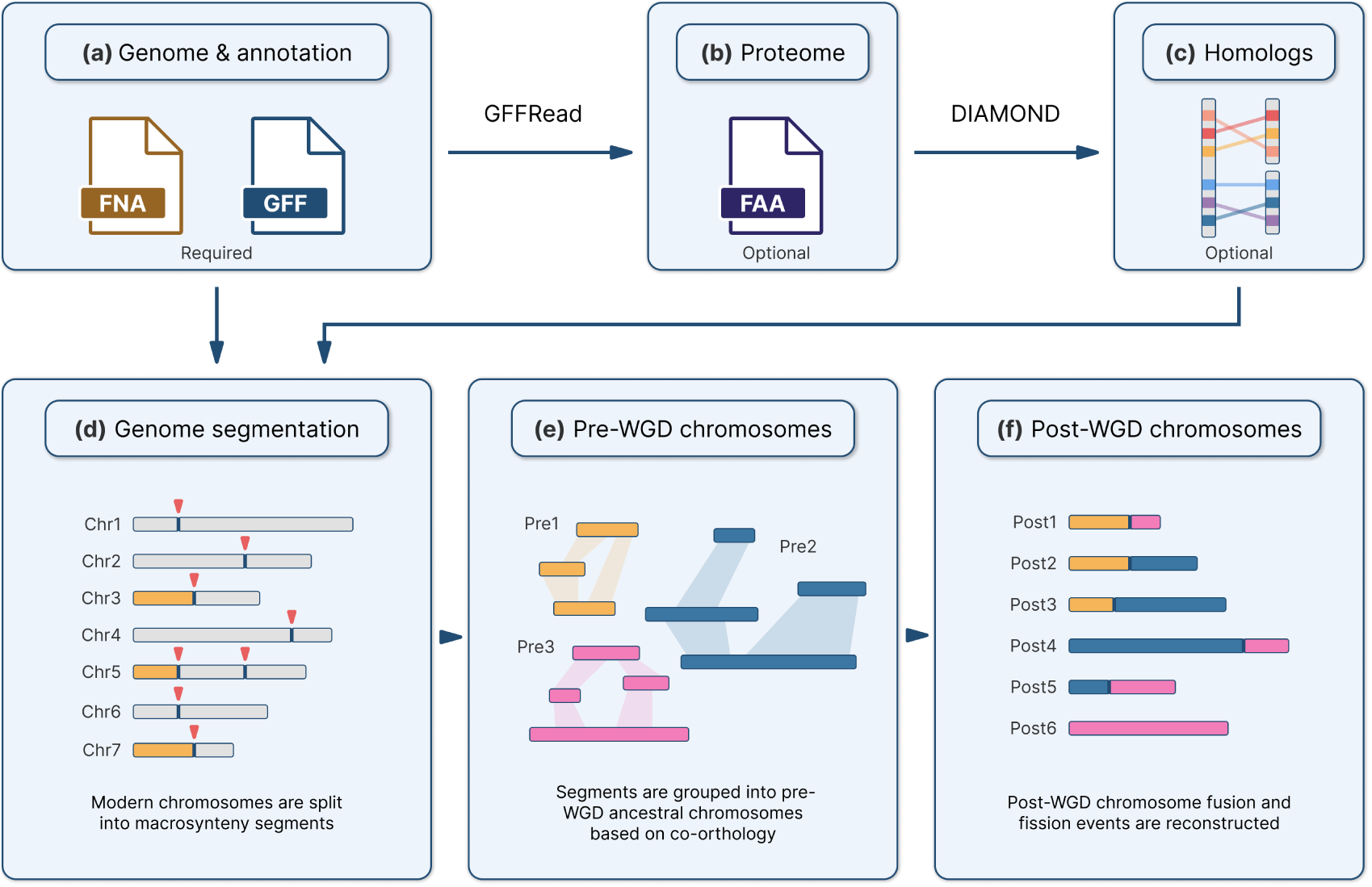
A flowchart highlighting the key steps in our WGD reconstruction pipeline. Required and optional inputs are annotated in panels a)-c). **a)** The minimum input required by our pipeline consists of genome sequence and annotation files. **b)** These files are then used to generate a proteome. Alternatively a proteome FASTA can be provided by the researcher. **c)** We perform a sequence similarity search to identify one-to-*n* homologs based on the number of WGD events separating the query and subject species. For example, we identify one-to-two orthologs between spotted gar and sturgeon as these genomes are separated by a single WGD event, resulting in additional duplicate copies in sturgeon (see ‘Methods’). **d)** We identify genomic segments that originated from a single ancestral chromosome using our sequence segmentation algorithm (see ‘Methods’). Each segment represents a chromosomal region that has remained unbroken, *i.e.,* has not experienced internal chromosomal fissions or fusions, since the WGD event. **e)** We arrange genomic segments from multiple genomes into pre-WGD chromosomes based on their co-orthology, *i.e.,* we cluster segments that are highly enriched in orthologous relationships. Importantly, these chromosomes are closer to ‘bags of segments’ and are not contiguous. **f)** We divide each pre-WGD chromosome into multiple post-WGD chromosomes based on an initial ohnolog set from the homology inference step, arranging segments such that homologous chromosomes are enriched in ohnologs. Based on the pre-WGD and post-WGD reconstructions, we can infer any chromosomal rearrangement events that occurred in this time-frame.

The core stages of our pipeline rely on identifying genomic segments that have remained unbroken, *i.e.,* have not experienced internal chromosomal fissions or fusions, after WGD. These segments are then arranged into ancestral linkage groups or ‘chromosomes’ at different evolutionary time-points, for example before and after WGD. As such, we developed a Bayesian segmentation algorithm to quickly and accurately delineate these segments (Fig. 1d, see ‘Methods’). Previously, this has been accomplished using chromosome alignments (Armstrong et al., 2020) or microsynteny-based analyses (Proost et al., 2012; Singh and Isambert, 2019), however, these approaches tend to be time consuming and not well suited to highly divergent genomes (Braasch et al., 2016). Instead, our pipeline relies on macrosynteny, and considers the conservation of gene content rather than gene order between formerly duplicated genomic regions (Nakatani and McLysaght, 2017; Simakov et al., 2020; Nakatani et al., 2021). This approach has been shown to consistently perform well across a wide range of taxa and divergence times (Nakatani and McLysaght, 2017; Simakov et al., 2020; Nakatani et al., 2021; Simakov et al., 2022).

Following this, the reconstruction of pre-WGD and post-WGD chromosomes is based on the enrichment or depletion of co-orthology and co-paralogy between previously identified genomic segments (Fig. 1e & f). For example, we might assign a group of segments as having originated from the same pre-WGD chromosome if they are enriched in both orthologs and paralogs. Conversely, there should be few paralogous relationships between segments originating from a single post-WGD chromosome (see ‘Methods’). This approach is flexible meaning that a) it can be applied to lineages with consecutive WGDs (Nakatani et al., 2021); and b) by changing the scoring function to determine an alternative ‘best’ arrangement of segments into chromosomes, it can be adapted to perform reconstructions that may not rely on WGD (see ‘Methods’). Finally, using reconstructed chromosomes we can infer fusion and fission events by investigating the enrichment of homologs between pre-WGD and post-WGD chromosomes. This can be informative of the relative timing of these events, *i.e.,* whether they happened before or after duplication, as well as their frequency, thus providing insight into macro-evolutionary processes shared by all descendant lineages.

One central goal of this work was to make the tool user-friendly. We have organised these steps into independent pipeline stages with the help of Snakemake (Mölder et al., 2021) which confers a number of benefits: Firstly, input can be provided at any point in the pipeline. If a researcher wishes to use a custom dataset, for example, a high-quality ohnolog dataset required for the reconstruction steps, the pipeline can be easily configured to use this instead of generating one *ad hoc*. Similarly, the pipeline can be freely stopped and restarted at any point. Output from one stage can be post-processed using custom scripts before passing it onto the next stage. Such options might be preferred by the expert user, though we have chosen sensible defaults to make this an ‘out of the box’ experience for a wide range of datasets. The behaviour of these stages can also be adjusted using a global configuration file. This not only makes it very flexible, but also allows the pipeline to determine the required input files for each stage, relaying this information to the user. Furthermore, since inputs can be determined ahead of time, work can be split into individual self-contained tasks that can be run in parallel, making them suitable for high-performance computing (HPC) clusters.

### Reconstructing the pre-WGD and post-WGD acipenseriform genomes

Together, sturgeons and the american paddlefish represent the extant Acipenseriformes, a lineage of non-teleost ray-finned fish. Having experienced an ancestral WGD event around 250 million years ago (Redmond et al., 2023), this clade has recently seen interest as a model for evolution after WGD, in no small part due to their slowly evolving genomes and recent availability of high-quality reference genome assemblies (Du et al., 2020; Cheng et al., 2021). In particular, recent work by Redmond et al. (2023) has cemented paddlefish and sturgeon as a model of asynchronous rediploidisation (Robertson et al., 2017; Redmond et al., 2023), a process of broad interest that may have impacted all vertebrate genomes (Marĺetaz et al., 2024; Yu et al., 2024), as well as those of many other paleopolyploid lineages. Reconstructing the ancestral acipenseriform genome has the potential to augment studies of this WGD, including by furthering our understanding of macroevolutionary processes underlying asynchronous rediploidisation, making it an excellent case study to demonstrate the efficacy of our pipeline. While this may not be the most technically difficult test-case, such as the highly shattered genomes of non-marine annelids (Lewin et al., 2024; Vargas-Chávez et al., 2025), or genomes that experienced older and/or multiple consecutive duplications, such as 2R in vertebrates (McLysaght et al., 2002; Dehal and Boore, 2005; Nakatani et al., 2021), this reconstruction demonstrates the performance of our pipeline in the context of limited data: Here we are constrained to only two post-WGD genomes, providing a good indication of the expected performance in poorly sampled clades.

First, we segmented the american paddlefish and sterlet sturgeon genomes against each other and against 6 outgroup genomes (see ‘Methods’). We found that this number of segmentations provided a consistent set of genomic breakpoints, with any additional genomes resulting in little gain in breakpoint consistency at the cost of additional runtime (Table S1). This resulted in 135 and 121 segments, respectively, in paddlefish and sturgeon that have remained unbroken after WGD. While excessive rearrangements could make segment boundaries difficult to delineate, especially for small segments with similar ortholog distributions (Fig. 2a), these genomes had not experienced much internal gene shuffling or large-scale chromosomal rearrangements, making the locations of most break-points easy to identify and within the detection thresholds of our segmentation model (Fig. 2b). Next, we partitioned these segments into 31 ancestral pre-WGD chromosomes using spotted gar as an outgroup (Fig. 2c, also see Table S2). This number is similar to chromosome counts observed in outgroup species, although these vary from 18 in bichir (Bi et al., 2021) to 29 in spotted gar (Braasch et al., 2016). Conversely, chromosome counts appear highly conserved in acipenseriformes, with higher counts in certain sturgeon species often appearing in multiples, directly linking them to consecutive rounds of WGD (Ludwig et al., 2001; Fontana et al., 2007; Havelka et al., 2011; Rajkov et al., 2014; Trifonov et al., 2016; Du et al., 2020; Cheng et al., 2021; Redmond et al., 2023). Given the large evolutionary time-frames and chromosomal plasticity of ray-finned fish, chromosome counts in outgroup genomes provide only approximate bounds for the expected chromosome count in the acipenseriform ancestor. Nonetheless, the 31 pre-WGD chromosomes that we infer are not inconsistent with these broad expectations, and closely match previous predictions that the pre-WGD acipenseriform ancestor had around 30 chromosomes (Birstein et al., 1997; Fontana et al., 2007; Trifonov et al., 2016).

**Fig. 2.**
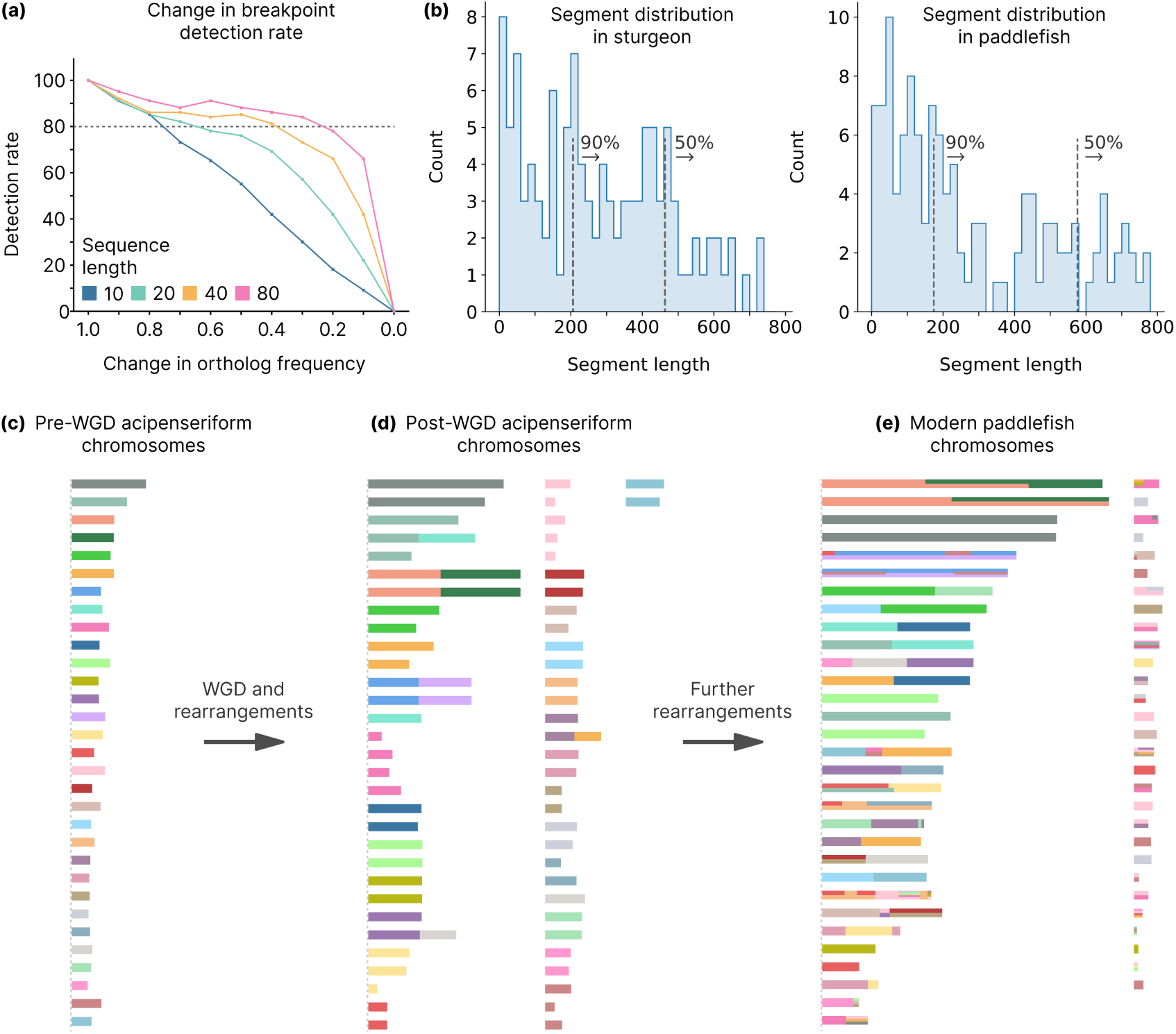
Using our pipeline to reconstruct the history of acipenseriform chromosomal evolution. **a)** We investigated the ability of our sequence segmentation algorithm to detect simulated known breakpoints (y-axis) given different combinations of ortholog frequency changes between genomic segments (x-axis, see ‘Methods’) and the total length of sequences that were segmented (coloured lines). Our model performs better, *i.e.,* correctly detects known breakpoints more often, with larger changes in ortholog frequency and increased sequence length. **b)** The distribution of inferred segment lengths in sturgeon and paddlefish, represented by the number of genes contained within each segment. Segments longer than 206 and 174 genes account for 90% of the genes in sturgeon and paddlefish respectively. Similarly, segments longer than 463 and 576 genes account for 50% of the genes in these genomes. **c)**-**e)** WGD events and major chromosome rearrangements in acipenseriform evolution are represented in chronological order from left to right. Each set of bars represents chromosomes at different stages of acipenseriform evolution. Colours indicate the relationship of segments to the pre-WGD chromosome reconstruction. These graphics were automatically generated as part of the pipeline, followed by manual curation (Supplementary Note 1). **c)** The ancestral acipenseriform had 31 chromosomes. **d)** Following whole genome duplication and chromosomal fusion and fission events, this number increased to 64. **e)** Finally, through additional rearrangement events, the modern paddlefish genome now has 60 chromosomes. The plot shows 62 scaffolds based on the annotation from Casey et al. (submitted) with filtering steps applied to remove small scaffolds with few genes (see ‘Methods’). Some chromosomal segments experienced irreversible mixing and their origin could not be traced back to a single ancestral chromosome. This shared evolutionary history is represented by coloured horizontal bars on paddlefish chromosomes.

We wanted to investigate whether this number is accurate and not an artefact caused by our choice of outgroups, genome quality, or segmentation settings. Our reconstruction is consistent when using spotted gar, bichir, or coelacanth as an outgroup. Using shark genomes resulted in a reconstruction with 25 pre-WGD chromosomes, although this is expected given the high degree of divergence between sharks and acipenseriformes compared to other, more suitable outgroups. A high degree of divergence makes irreversible rearrangements in either lineage more likely, obscuring comparisons between these genomes and has been known to lead to underestimating the number of ancestral chromosomes using our approach (Nakatani et al., 2021). To test whether genome quality or poor segmentations have an effect on inferred chromosome count, we introduced random genomic breakpoints to spotted gar, paddlefish, and sturgeon chromosomes. These should be analogous to fragmented assemblies and erroneous breakpoints. We found that our reconstruction was robust to the addition of up to 15% random breakpoints (Table S3), with larger increases having a significant effect on inferred chromosome count. Given that over 90% of our previously identified breakpoints were consistent across all segmentations (Table S1), this suggests that erroneous breakpoints are unlikely to have biased the ancestral chromosome count. Furthermore, this demonstrates that our pipeline and pre-WGD reconstruction are robust to some degree of noise introduced by low-quality genomes.

Finally, we partitioned pre-WGD segments into 72 post-WGD chromosomes and used these to infer the chromosomal fusion and fission events that occurred between these two states. However, the number of post-WGD chromosomes was likely inflated due to delayed and asynchronous rediploidisation in acipenseriformes, which is not modeled in our WGD reconstruction algorithm (Supplementary Note 1). This inflation does not compromise the overall reconstruction as most post-WGD chromosomes appear to be correctly resolved, *i.e.,* are inferred to be present in two copies, even though many of them rediploidised after the divergence of paddlefish and sturgeon (Redmond et al., 2023). Instead, the algorithm appears to be inconsistent in cases where chromosomes were partially rediploidised at the time of speciation, as was the case for the large acipenseriform chromosomes (Redmond et al., 2023), or possibly in cases where delayed rediploidisation is combined with multiple fusion or fission events. Fortunately, these inconsistencies are easily identified by comparing the post-WGD reconstruction to extant paddlefish and sturgeon chromosomes. For example, pre-WGD chromosomes 3 and 4 were originally inferred to have split into 4 post-WGD chromosomes (Figure S1). The fact that these chromosomes have remained intact in both paddlefish and sturgeon (Figure S2 & S3) suggests that this is likely an artefact rather than multiple fissions followed by independent fusions. After manual curation to correct these artefacts, we inferred that the ancestral acipenseriform likely had 64 chromosomes (Fig. 2c-e). Using scripts provided with the pipeline, we visualised the chromosome-level evolution of the acipenseriform genome at three different stages: Pre-WGD, post-WGD, and the extant paddlefish and sturgeon (Fig. 2c-e, Figure S2 & S3).

### Investigating synteny conservation with non-reconstruction outgroup genomes

Our reconstruction provides good coverage of the paddlefish and sturgeon genomes, including 99.6% of the genes that have been assigned to chromosomes in both species. As a validation of our reconstruction, we examined the macrosynteny conservation with representative outgroup genomes including grey bichir, coelacanth, and whale shark (Fig. 3, see ‘Methods’). Despite periods of major chromosomal rearrangements in vertebrate history, such as those caused by genomic instability following WGD (Nakatani et al., 2021), we can still observe strong conservation of macrosynteny between various vertebrate chromosomes, including in acipenseriformes (Venkatesh et al., 2014; Braasch et al., 2016; Du et al., 2020; Cheng et al., 2021; Nakatani et al., 2021). As such, we expect a reliable reconstruction to show a highly non-random distribution of orthologs in genomes that have not been used in the reconstruction. Indeed, we find that orthologs are not randomly scattered throughout modern genomes, but are clustered into a small number of chromosomes. Importantly, all of our reconstructed chromosomes map onto at least one chromosome in an outgroup genome, some of which exhibit one-to-many relationships which may be indicative of fusion or fission events. This is expected to some degree given large evolutionary time-scales (Inoue et al., 2010). Alternatively, this could indicate extensive chromosome mixing which could lead us to underestimate the number of ancestral acipenseriform chromosomes (Nakatani et al., 2021). However, most pre-WGD chromosomes map onto a single outgroup chromosome in at least one of our comparisons, suggesting that these one-to-many relationships are likely to reflect genuine lineage-specific rearrangements.

**Fig. 3.**
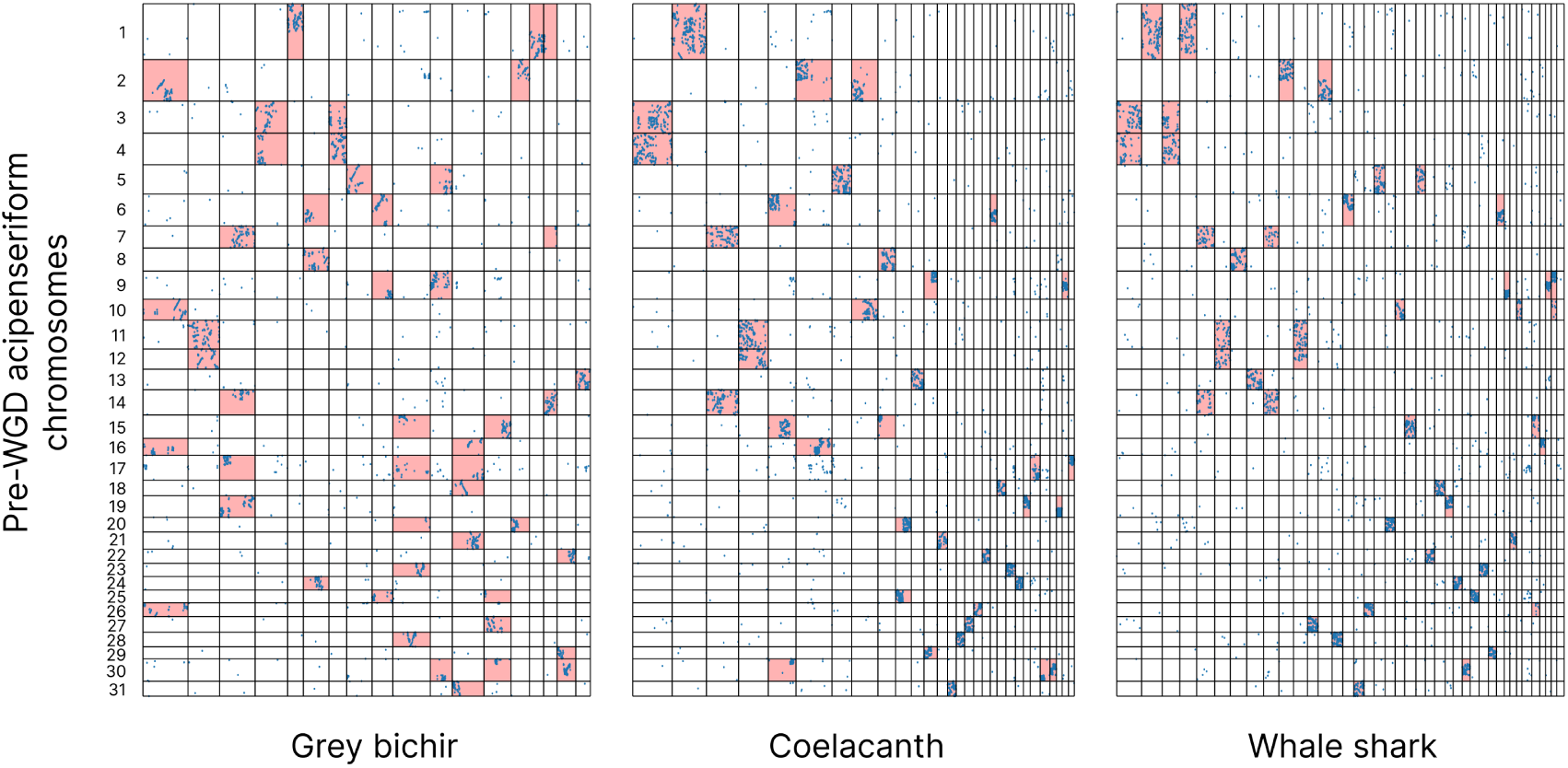
Each of the reconstructed pre-WGD chromosomes exhibits a distinct ortholog distribution in outgroup genomes. The reconstructed pre-WGD acipenseriform chromosomes, as represented by spotted gar segments (y-axis), were compared against grey bichir, coelacanth, and whale shark chromosomes (x-axis), all of which were not used in the reconstruction. The acipenseriform chromosomes (1-31) are shown from top to bottom along the y-axis, and their boundaries are indicated by horizontal black lines. Similarly, the bichir, coelacanth, and whale shark chromosomes are delineated by vertical black lines. The locations of orthologous genes between pre-WGD chromosomes and each of the three outgroup genomes were plotted and are indicated by blue dots. The distribution of these genes is highly non-random; clusters of orthologs are confined to distinct regions of each genome. Chromosome pairs for which this enrichment is significant are highlighted in red (see ‘Methods’).

Finally, our reconstruction appears to be sensitive enough to recover many small chromosomes, although size estimates are based on gene content and do not account for changes in the amount of non-coding DNA. Nevertheless, our reconstruction can be a good starting point to make inferences about macro-evolution in acipenseriformes, whose genomes contain both macro and microchromosomes. However, unlike in birds, there is no clear cut-off between large and small chromosomes in acipenseriformes, the impact of which is currently not well understood (Symonová et al., 2017; Du et al., 2020; Cheng et al., 2021; Huang et al., 2023). Based on macrosynteny conservation, we find that small chromosomes in sturgeon and paddlefish often correspond to small chromosomes in some outgroups, such as coelacanth and whale shark (Figure S4), both of which have microchromosomes (Amemiya et al., 2013; Kawaguchi et al., 2026). Furthermore, if we use reconstructed pre-WGD chromosomes as a proxy for ancestral chromosome size, we find that pre-WGD chromosome size generally corresponds to chromosome size in modern paddlefish and sturgeon genomes (Figure S5). Given this, we can infer that at least some microchromosomes in acipenseriformes likely pre-date the WGD. However, given our reliance on gene content to estimate chromosome size, we cannot be certain whether the unique distribution of chromosome size in acipenseriformes pre-dates the shared WGD, or is a direct consequence of it, as may have been the case with vertebrate microchromosomes (Nakatani et al., 2021).

Genome reconstruction tools have been instrumental in solving difficult evolutionary problems (Nakatani et al., 2021; Simakov et al., 2022), however, many such tools are resource and time intensive. While our pipeline is not directly comparable to most existing tools, as these have been designed to tackle a different set of problems, often involving microsynteny and short diveregence times (Proost et al., 2012; Muffato et al., 2023; Bernard et al., 2025), nevertheless, we have focused on optimisations aimed at reducing resource usage and total runtime. Our acipenseriform reconstruction ran to completion in less than 2 hours with 10 concurrent threads. Although this would take longer given a larger dataset, our pipeline should be suitable for use with consumer hardware; our pipeline can be used to quickly and easily reconstruct ancestral genomes with a high degree of coverage.

## Discussion

Here we present a pipeline for reconstructing ancestral WGDs. Previously, this has been done using disparate methods (Sacerdot et al., 2018; Simakov et al., 2020; Nakatani et al., 2021; Simakov et al., 2022) and is often an involved and time-consuming process. Our pipeline aims to solve this while being easy to configure and easy to use across a wide range of genomes and duplication events. Making ancestral reconstruction easily accessible has many advantages; WGD leaves traces in the form of synteny, which can serve as a valuable source of genomic information, whose potential has not been fully realised in previous work (Wolfe and Shields, 1997; McLysaght et al., 2002; Macqueen and Johnston, 2014; Nakatani et al., 2021; Redmond et al., 2023). Importantly, it can act as an alternative or supplementary source of information when other methods, such as phylogenetics-based approaches, are less viable. This is often the case in hard-to-solve problems involving ancient WGD with high degrees of ohnolog loss, consecutive WGD (Nakatani et al., 2021), or in the face of systematic bias, such as large differences in nucleotide/amino acid composition between genomes (Fried et al., 2003; Yu et al., 2024; Marĺetaz et al., 2024). Furthermore, this approach allows for reconstructions without an outgroup, which expands the potential use cases to poorly sampled parts of the tree of life. Since reconstructions allow us to identify homologous segments, and thus genes duplicated by WGD, they could be used as an alternative approach to root gene trees on duplication nodes in the absence of a suitable outgroup (Emms and Kelly, 2017), although this would likely be complicated by delayed rediploidisation (Supplementary Note 1). This may be helpful in extending ohnolog datasets to include gene families where an outgroup gene has been lost or cannot be easily identified, or in solving difficult cases such as resolving the number and timing of WGD event after whole genome shattering in non-marine annelids (Vargas-Chávez et al., 2025). Of course, phylogenetic analysis can be used to inform the reconstruction, for example, by specifying homolog sets based on Ensembl Compara (Yates et al., 2019), as has been done previously (Nakatani et al., 2021). Similarly, a highly curated ohnolog set can be provided as input to improve the accuracy of the reconstruction instead of inferring one *ad hoc* (Niezabitowski et al., forthcoming). Our Bayesian sequence segmentation algorithm is a core component of the pipeline. Unlike many models that require Markov Chain analysis to estimate the posterior (Gilks, 1996), our model is mathematically tractable and thus can be computed in a deterministic manner, making it simple, predictable, and suitable for consumer hardware. Our high-performance Rust implementation, specifically created for use in the pipeline, uses predetermined default settings that should be suitable for a wide range of orthology-based sequence segmentation tasks. Specifically, our model is highly reliable at identifying medium and large genomic segments. It also performs reasonably well at identifying very short segments given a high enough change in ortholog frequency. Furthermore, our Rust implementation has a low rate of false-positives. Since these errors are by definition random, the false-positive rate can be further reduced by performing multiple segmentations followed by postprocessing steps such as our sliding window analysis. This may not be possible when looking for lineage-specific rearrangements or in the case of poorly sampled clades, or clades with few extant species (Yu et al., 2024; Marĺetaz et al., 2024). In this case, breakpoints that fall outside the acceptable model performance thresholds can be removed entirely. Finally, while the Rust version of our model is intended for large generic datasets, we also provide a reference implementation in Python that be can easily finetuned and adjusted for more powerful inference in smaller, more specialised problems. In short, our model is accurate and adaptable to a wide range of tasks.

Our pipeline performs well in reconstructing the ancestral acipenseriform genome, with good coverage of paddlefish and sturgeon genes, and providing a near perfect match to the long standing prediction that this genome would have 30 chromosome pairs (Birstein et al., 1997; Fontana et al., 2007; Trifonov et al., 2016). Given the chromosome counts in the most closely related lineages (Braasch et al., 2016; Bi et al., 2021), the reconstructed ancestor of bony vertebrates (31) (Nakatani et al., 2007), and similar approaches used in reconstructing the vertebrate ancestor (Nakatani et al., 2021), our pre-WGD chromosome count should be accurate, assuming there was no extensive chromosome mixing in the acipenseriform ancestor. Indeed, there has been surprisingly little rearrangement after WGD in contrast to extensive shuffling after the vertebrate 2R (Simakov et al., 2020; Nakatani et al., 2021). This is despite the 450 million years of evolution separating some of the genomes in our comparison (Inoue et al., 2010). While our pipeline generally performs well, the post-WGD acipenseriform reconstruction highlights a limitation in our methodology. Because current probabilistic macrosynteny models assume a uniform paralog distribution after WGD, they are highly sensitive to the heterogeneous blocks of paralogs generated by asynchronous rediploidisation (Nakatani et al., 2021). Consequently, there is a need for future reconstruction algorithms to explicitly integrate rediploidisation timing as a parameter. Incorporating both the spatial conservation and temporal resolution of ohnologs would not only prevent the artefacts observed in our reconstruction, but could also automate the identification of differentially rediploidising genomic regions. Nevertheless, our reconstruction can contribute to the ongoing work in understanding rediploidisation (Redmond et al., 2023). This process entails the return of tetraploid chromosomes to a more stable diploid state (Wolfe, 2001; Conant and Wolfe, 2008; Robertson et al., 2017; Mandáková and Lysak, 2018; Du et al., 2020; Redmond et al., 2023) and is thought to occur through rearrangements and other mutations (Robertson et al., 2017; Redmond et al., 2023; Xie et al., 2026). Ancestral genome reconstructions provide the necessary framework to verify these mechanisms, as recently demonstrated by work utilising our dataset to link shifts in rediploidisation timing to chromosomal rearrangement events in acipenseriformes (Casey et al., submitted).

Our pipeline improves on previous work by providing a standardised and user-friendly way to reconstruct ancestral WGD events, allowing researchers to better exploit the syntenic traces that remain scattered throughout the genome. As we have shown with our reconstruction of the acipenseriform WGD, this can allow us to gain better insight into various genomic processes and contribute to multiple lines of research. We expect that the ease of installation and ease of use of this pipeline will provide the opportunity for a greater number of researchers to analyse their genomes of interest and make novel and interesting discoveries.

## Materials and Methods

### Homology inference

We used DIAMOND (Buchfink et al., 2021) to perform a sequence similarity search for homology identification. For each query gene, we defined its orthologs as the top *n* best scoring DIAMOND hits, where *n* is based on the number of WGDs separating the query and subject species (Table S4). For example, we found 1:1 orthologs between paddlefish and sturgeon as their genomes share the same number of WGD events. However, compared to bichir, the paddlefish genome has experienced 1 additional WGD event. As a result, we found 1:2 orthologs between these genomes.

We generated an ohnolog set required for our ancestral reconstructions by identifying paralogs whose duplication timing coincides with that of the acipenseriform WGD event. Given two reference species, a pair of genes with a bitscore *s* was annotated as a paralog pair if they met the following conditions: a) The shorter gene does not have a larger than *s* bitscore to any genes from genome *A*. b) The shorter gene has a larger than *s* bitscore to the best-matching gene from genome *B*. Genome trios for this analysis are shown in Table S5. In order to exclude large gene families, we discarded the gene pair if it was not one of the top *n* best scoring DIAMOND hits. The value of *n* was determined by the number of expected duplicates given the WGD history of a species (Table S4). In addition, paralogs located within 5 genes of each other on the same chromosome were discarded.

### Genome segmentation

We developed a Bayesian sequence segmentation algorithm in order to identify genomic regions of conserved macrosynteny (Auger and Lawrence, 1989; Liu and Lawrence, 1999; Nakatani and McLysaght, 2017). Macrosynteny can be defined as the conservation of gene content between homologous chromosomal regions (Fig. 4a & b). We can identify the boundaries (hereafter breakpoints) separating these regions on some chromosome Q, by identifying changes in the frequency of homologs along the chromosome when compared to some other chromosome S. In brief, the model works as follows: Let *R* be the observed sequence of genes along chromosome Q. These genes are split into two categories *c*, based on their similarity to genes on some other chromosome S; *c* = 0 in the absence of any similarity and *c* = 1 if the gene has at least one homolog on S. Given *κ*, the number of synteny breakpoints in sequence *R*, the probability of observing such a sequence is given as follows:

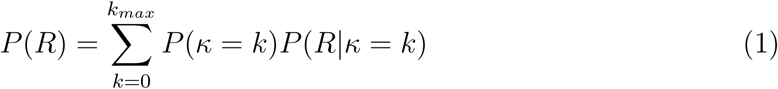

**Fig. 4.**
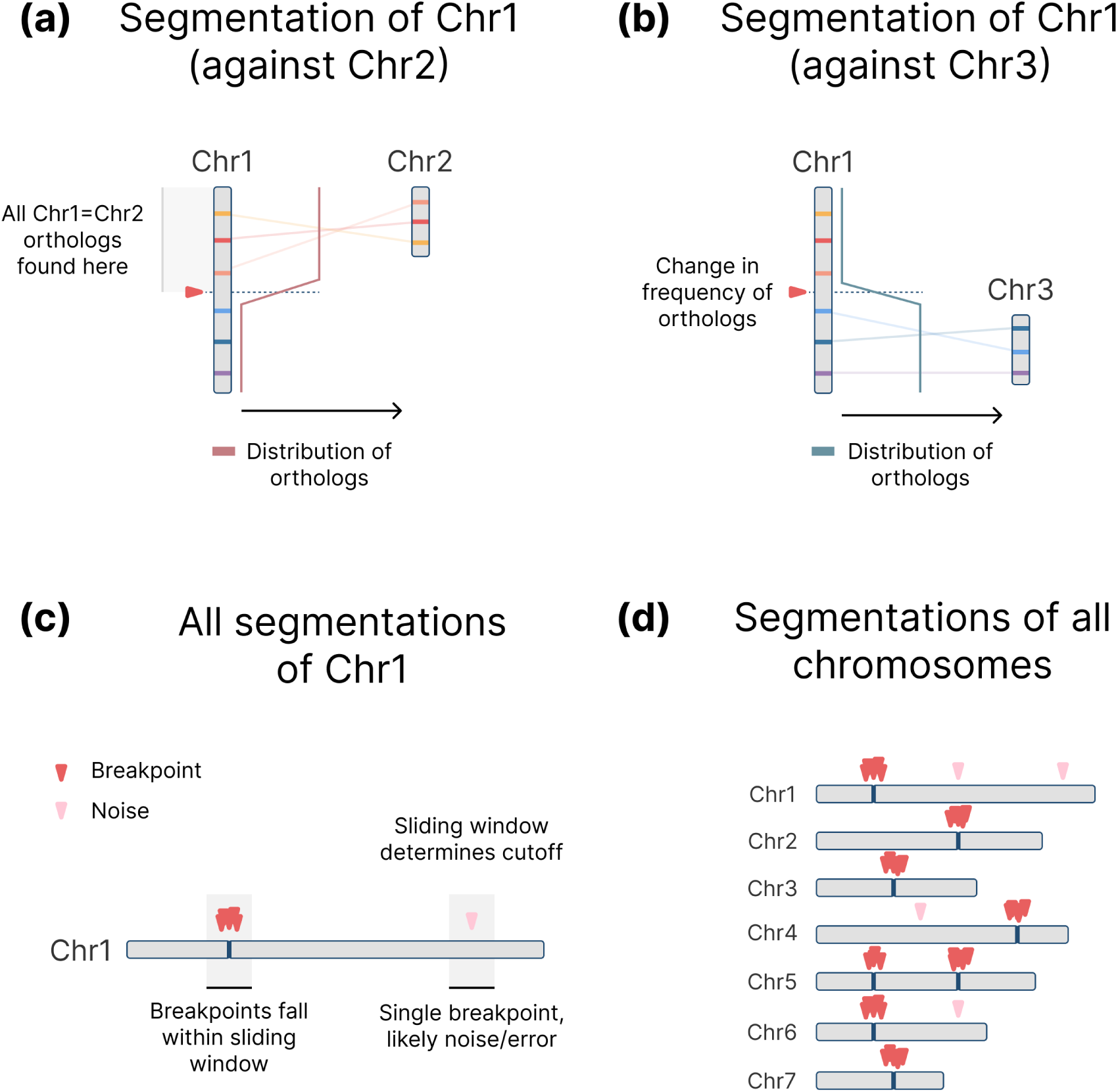
Genome segmentation algorithm. **a)** A hypothetical segmentation of chromosome 1 against some chromosome 2 from another genome. We identify the orthologs between the two chromosomes and plot their frequency along chromosome 1. The position on chromosome 1 at which this frequency changes is a breakpoint, delineating the location of a chromosomal fission or fusion event. **b)** We can repeat this process, this time segmenting chromosome 1 against chromosome 3. **c)** Repeating this process, we can segment chromosome 1 against all other genomes in our dataset. This allows us to identify false-positives or lineage-specific breakpoints that are not useful in our analysis. We can use a sliding window analysis to determine if a breakpoint should be kept based on its proximity to other breakpoints; it is unlikely that false-positives fall in the same location over multiple segmentations. **d)** This process is repeated for all the chromosomes and genomes in our dataset.

We can then calculate *P* (*κ* = *k*)*P* (*R|κ* = *k*) given the formula below, where *A* represents the segmentation of sequence *R*, referring to the number and position of synteny breakpoints, *n_k_* is the number of genes in segment *k*, *n_k,c_* is the number of genes in segment *k* that belong to category *c*, and *α_c_* is the prior for category *c*:

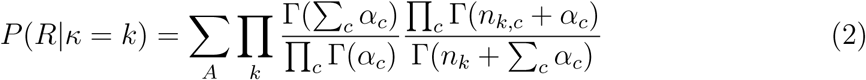

This allows us to calculate the number of breakpoints in the chromosome up to some value *k_max_*. Since the model is effectively intractable for values of *k >* 2 using a brute-force approach, we used dynamic programming to complete the calculation. Once we know the number of breakpoints in a chromosome, we can work backwards in order to find the locations of these breakpoints. A detailed explanation and a reference implementation in Python can be found on GitHub (github.com/McLysaght-Evolutionary-Genetics/bayesian-segmentation). The implementation used in our analyses was written in Rust and called using Python FFI, significantly increasing the performance of our pipeline.

In order to test the detection rate of our model under various conditions, we generated test sequences, varying the length and change in ortholog frequency between segments (Table S6). Similarly we tested the false-positive rate by checking if our model detected any breakpoints in sequences with a fixed ortholog frequency (Table S7). We generated 1000 test sequences for each combination of frequency and sequence length. Low sequence lengths and, in the case of detection rate, small changes in frequency represent the worst case scenario for our model.

We used the following parameters for all comparisons: *k* = 20, *α* = 1.0, *β* = 1.0. For the acipenseriform reconstruction, we segmented the american paddlefish (*Polyodon spathula*) and sterlet sturgeon (*Acipenser ruthenus*) genomes against each other and against spotted gar, grey bichir, coelacanth, elephant shark, whale shark, zebra shark, and little skate. Post-processing steps were applied to the resulting breakpoints to remove any false-positives as well as any lineage-specific breakpoints: Breakpoints were kept if at least *n* breakpoints appeared within a sliding-window of 5 genes. The value of *n* is equal to the number of genomes used in the segmentation (Fig. 4c & d).

### Ancestral genome reconstruction

A modified version of the reconstruction algorithm from Nakatani et al. (2021) was used to reconstruct the ancestral acipenseriform genome (Supplementary Note 2). We used spotted gar, american paddlefish, and sterlet sturgeon for the reconstruction. The spotted gar genome was used as a reference. Default parameters were used unless specified otherwise. The reconstruction was performed with values of *K* ranging from 20-80 and the most significant reconstruction was selected using a hypergeometric test. This test compares the distribution of all possible gene pairings to the distribution of observed paralog pairs in order to detect the enrichment of paralogs within pre-WGD chromosomes. Let *g* and *p* be the number of gene and paralog pairs in the genome respectively. Let *v* be the number of gene pairs that both derive from the same pre-WGD chromosome. Let *X* be the number of paralog pairs that both derive from the same pre-WGD chromosome and *x* be the observed number of such pairs. The significance of reconstruction, *x*, is given as follows:

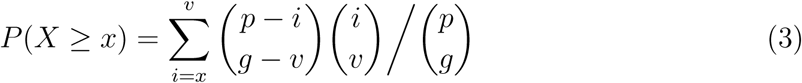

We identified chromosomal rearrangements between pre-WGD and post-WGD chromosomes by comparing the enrichment of orthologs between these two sets as per Nakatani et al. (2021). To identify ancestral chromosome pairs that share a significant number of orthologs, we used the Poisson distribution, assuming that orthologs are randomly distributed. Relationships were classified as significant where *P <* 10*^−^*^2^. Rearrangements were classified as having occurred before the WGD event if at least two post-WGD chromosomes shared synteny with two or more pre-WGD chromosomes. Otherwise, we assumed that the rearrangement occurred after the WGD event. A similar approach was used in Fig. 3 to identify ortholog enrichment between pre-WGD and modern chromosomes. Pre-WGD chromosome lengths in Fig. 2 were defined as the numbers of spotted gar genes assigned to each reconstructed chromosome. The lengths of post-WGD chromosomes were defined by 2*x*, where *x* is the number of post-WGD genes retained from each pre-WGD chromosome. A pre-WGD gene was inferred to be retained on the post-WGD chromosome if the gene has at least one ortholog in paddlefish or sturgeon segments assigned to the post-WGD chromosome.

## Acknowledgments

We thank members of the Molecular Evolution research group for helpful discussion and comments.

## Funding

This work was supported by funding from the European Research Council, grant agreement 771419 (A.McL.).

## Author contributions

Conceptualisation: LN, AMcL, AKR

Data curation: LN

Formal analysis: LN

Funding acquisition: AMcL

Investigation: LN

Methodology: LN

Project administration: AMcL

Resources: AMcL

Software: LN

Supervision: AMcL

Validation: LN

Visualisation: LN

Writing – original draft: LN

Writing - review & editing: LN, AMcL, AKR

## Competing interests

The authors declare that they have no competing interests.

## Data and materials availability

All data are available in the main text or the supplementary materials. The reconstruction pipeline is available on GitHub (https://github.com/McLysaght-Evolutionary-Genetics/macrosynteny-toolkit/).

## Supplementary Materials

### The PDF file includes

Supplementary Notes 1 to 2 Figures S1 to S5

Legends for Figures S1 to S5 Tables S1 to S7

Legends for Tables S1 to S7

## Supplementary information for: An automated pipeline **for reconstructing whole genome duplications**

### Supplementary Note 1. The effect of delayed rediploidisation on the acipenseriform reconstruction

Following autopolyploidisation, duplicated chromosomes are initially identical and pair as multivalents during meiosis. The transition from this state back to stable, bivalent diploidy is a process known as rediploidisation. While this was traditionally assumed to occur rapidly and uniformly across the genome, recent evidence demonstrates that it can be delayed and proceed asynchronously over millions of years. This results in discrete genomic blocks resolving at different times (Robertson et al., 2017; Parey et al., 2022; Redmond et al., 2023; Xie et al., 2026). Structural rearrangements, such as inversions (Casey et al., submitted) or unbalanced fusions (Xie et al., 2026), are thought to locally suppress multivalent recombination, establishing distinct regions that rediploidise at dif-ferent time-points (Robertson et al., 2017; Gundappa et al., 2022). At the time of the paddlefish-sturgeon divergence, approximately 60% of the ancestral acipenseriform genome was still undergoing multivalent pairing (Redmond et al., 2023). Modern paddlefish and sturgeon genomes are therefore mosaics of regions that rediploidised before speciation (Ancestral Ohnolog Resolution; AORe) and regions that rediploidised independently after speciation (Lineage-specific Ohnolog Resolution; LORe) (Robertson et al., 2017; Redmond et al., 2023).

The probabilistic macrosynteny model employed in our pipeline (adapted from Nakatani et al. (2021)) relies on identifying a continuous, genome-wide distribution of paralogs to link pre-WGD and post-WGD chromosomes. This approach implicitly assumes that WGD was followed by rapid, uniform rediploidisation, resulting in paired ohnologous regions with a relatively homogeneous paralog distribution. Delayed and asynchronous rediploidisation inherently violates this assumption. Genomic regions that rediploidised after the paddlefish-sturgeon divergence (LORe) share exceedingly high sequence similarity. This similarity frequently causes these regions to either (i) suffer from assembly collapse in modern reference genomes, although this is resolved in our dataset by leveraging a duplicate-resolved paddlefish genome (Casey et al., submitted), or (ii) evade standard ortholog/paralog detection thresholds due to low sequence divergence and lineage-specific phylogenetic topologies (Redmond et al., 2023). When applied to a chromosome composed of regions with varying rediploidisation histories, the algorithm is likely to identify strong paralog signals in early-rediploidising (AORe) regions, followed by differences in paralog density at the boundaries of late-rediploidising (LORe) regions. These differences in signal may be interpreted as ancestral chromosome breakpoints, which can lead to artefactually fragmenting continuous chromosomes along the boundaries of these regions. This variation in paralog density likely explains the inflated post-WGD chromosome count initially output by our pipeline (72 post-WGD chromosomes). While the algorithm accurately reconstructed the majority of the genome following a standard 1:2 duplication ratio, several large pre-WGD macrochromosomes were artefactually fragmented into three, four, or even six post-WGD segments. Fortunately, these inconsistencies can be identified and corrected by mapping the fragmented post-WGD segments back to extant paddlefish and sturgeon genomes. One specific artefact we observed is a 1 to 4 to 2 pattern, where the algorithm infers that a single pre-WGD chromosome split into four post-WGD segments, yet comparative mapping reveals these four segments are placed contiguously on exactly two intact chromosomes in both modern paddlefish and sturgeon, for example, pre-WGD chromosomes 3 and 4 (Figure S1). Biologically, it is highly improbable that an ancestral chromosome underwent multiple fissions immediately after WGD, only for those exact segments to independently fuse back together in identical configurations in multiple descendant lineages. Rather, these regions likely represent ancestral tetravalents, which are chromosomes that were still recombining as single units at the time of acipenseriform speciation. To account for this, we performed manual curation on the output generated by our pipeline (Fig. 2). We merged inferred post-WGD chromosomes where a single pre-WGD chromosome was fragmented into multiple post-WGD segments that currently form intact, unbroken linkage groups in extant genomes. This curation should filter out the algorithmic noise caused by asynchronous rediploidisation, resolving the inflated 72-chromosome count to an ancestral post-WGD karyotype of 64 chromosomes.

### Supplementary Note 2. Modifications to the probabilistic macrosynteny model

The genome reconstruction software used in our pipeline is a modified version based on Nakatani et al. (2021). We removed several statistical utility functions and replaced them with more performant alternatives. Unlike the previous versions, these have permissive licenses and can be publicly shared without any workarounds. We also changed the program architecture to allow for larger datasets. Previously, attempting reconstructions with large numbers of genomic segments would result in a crash. Importantly, these changes did not have any effect on inference. The modified version of the algorithm is available on GitHub (https://github.com/McLysaght-Evolutionary-Genetics/ancestral-reconstruction).

---

**Figure S1:**
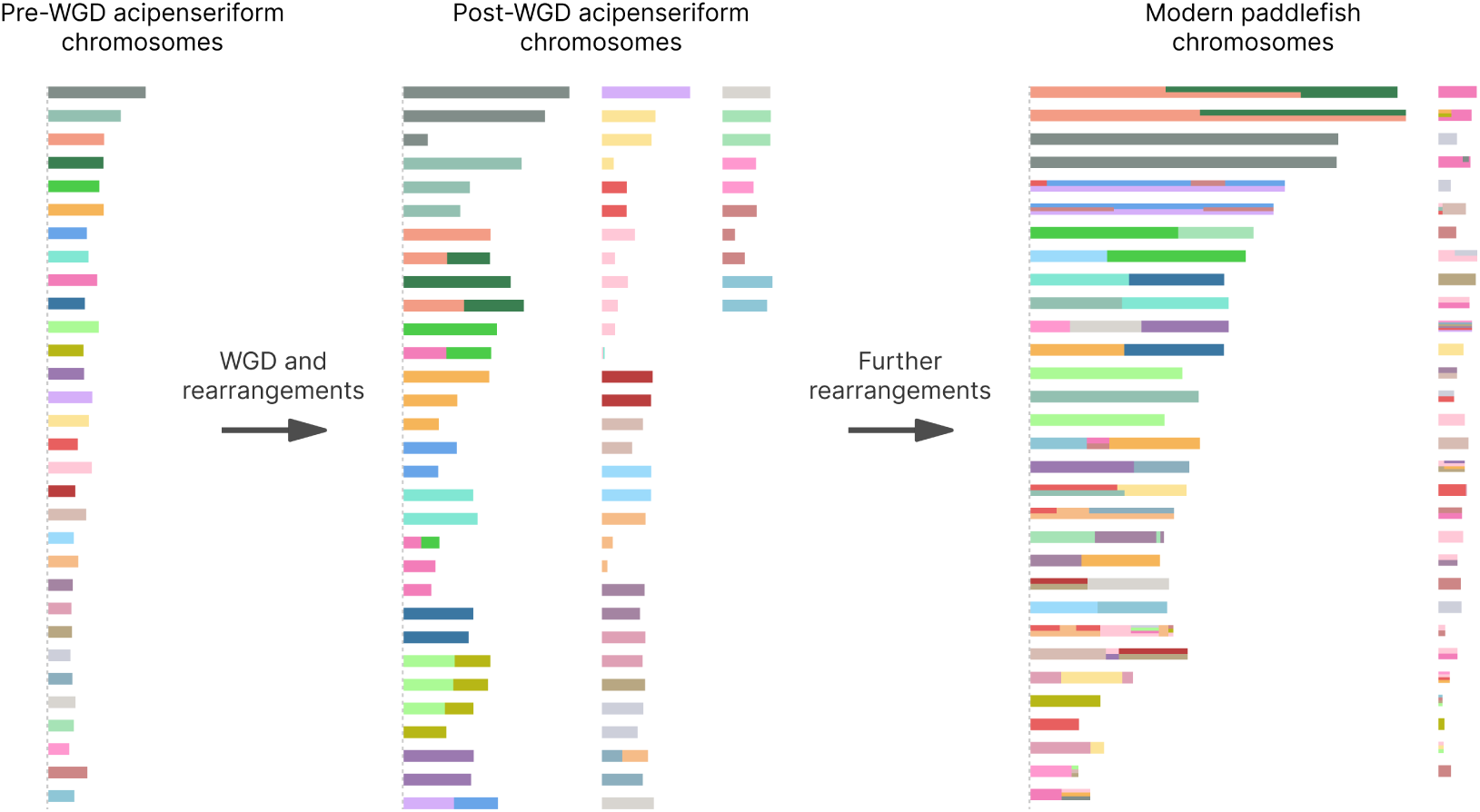
Model of chromosomal evolution in acipenseriformes prior to manual curation. WGD events and major chromosome rearrangements in acipenseriform evolution are represented in chronological order from left to right. Each set of bars represents chromosomes at different stages of acipenseriform evolution. Colours indicate the relationship to the pre-WGD chromosome reconstruction. The ancestral acipenseriform had 31 chromosomes. Following whole genome duplication and chromosomal fusion and fission events, this number increased to 72. This inflated count is likely an artefact caused by delayed and asynchronous rediploidisation in acipenseriformes and has been manually fixed in Fig. 2. Finally, through additional rearrangement events, the modern paddlefish genome now has 60 chromosomes. The plot shows 62 scaffolds based on the annotation from Casey et al. (submitted) with filtering steps applied to remove small scaffolds with few genes (see ‘Methods’). Some chromosomal segments experienced irreversible mixing and their origin could not be traced back to a single ancestral chromosome. This shared evolutionary history is represented by coloured horizontal bars on paddlefish chromosomes.

**Figure S2:**
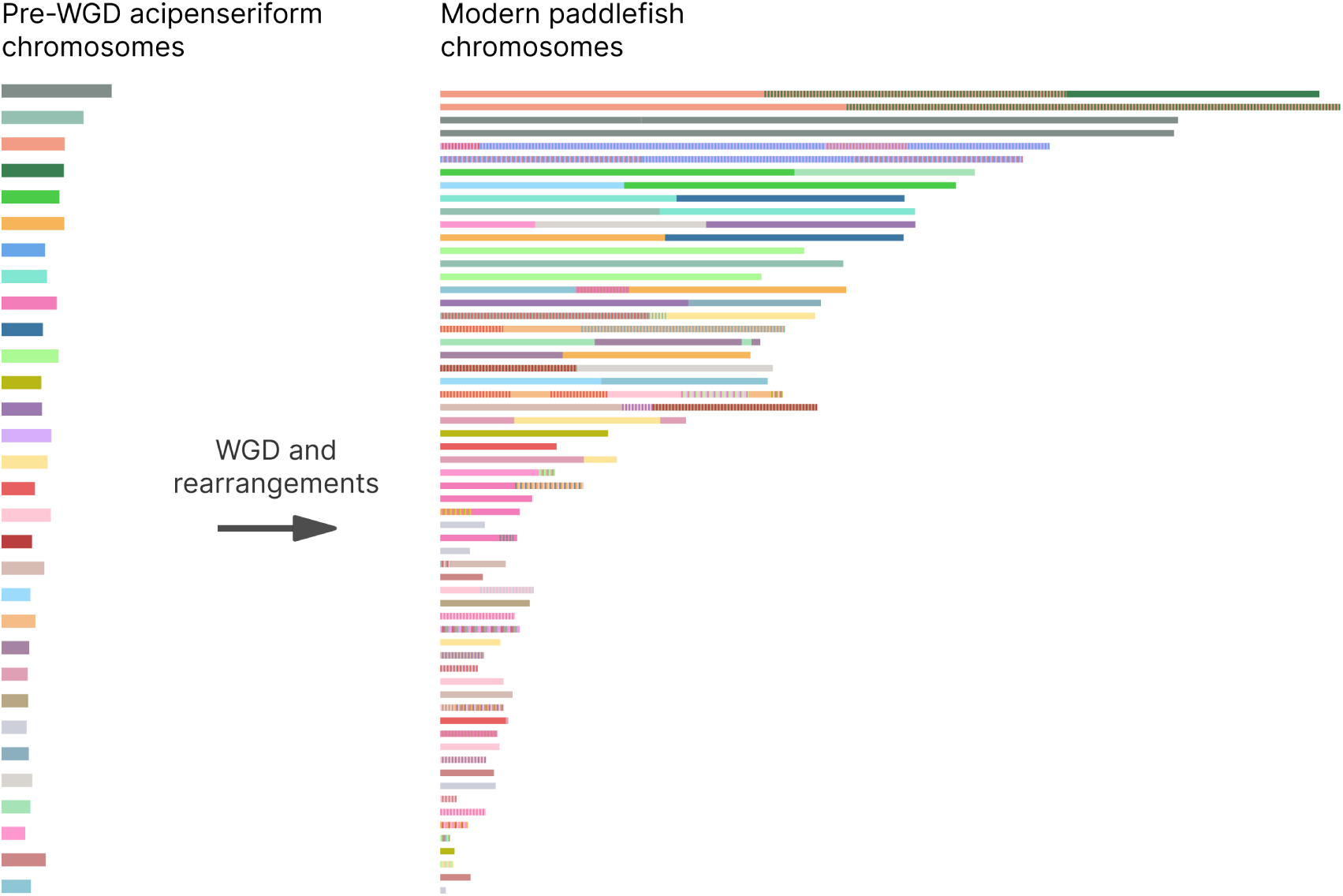
Modern paddlefish segments coloured based on their relationship to reconstructed pre-WGD chromosomes. Each set of bars represents chromosomes at different stages of acipenseriform evolution. The ancestral acipenseriform had 31 chromosomes. Through WGD, followed by chromosomal fusion, fission, and rearrangement events, the modern paddlefish genome now has 60 chromosomes. The plot shows 62 scaffolds based on the annotation from Casey et al. (submitted) with filtering steps applied to remove small scaffolds with few genes (see ‘Methods’). Some chromosomal segments experienced irreversible mixing and their origin could not be traced back to a single ancestral chromosome. This shared evolutionary history is represented by coloured stripes on paddlefish chromosomes.

**Figure S3:**
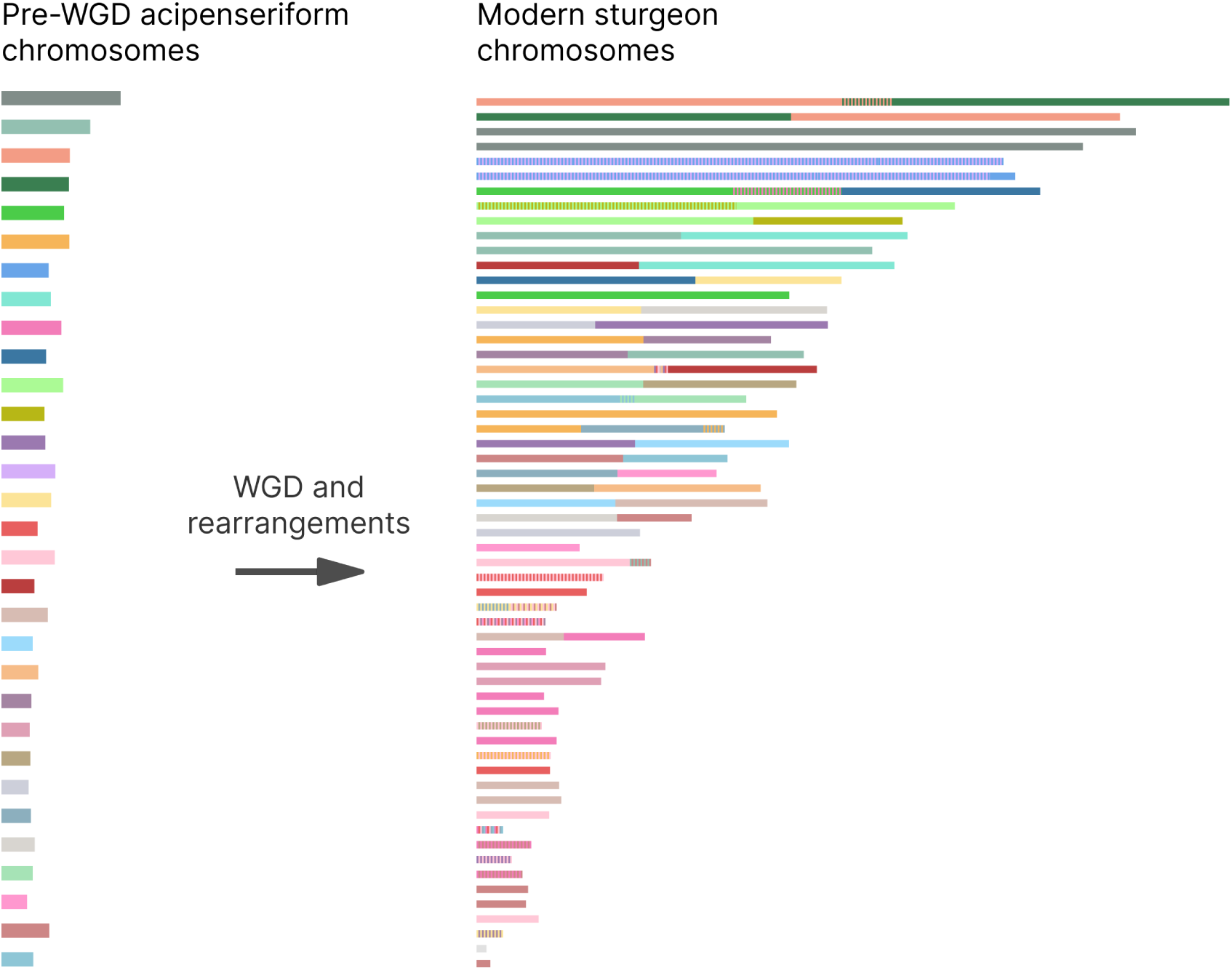
Modern sturgeon segments coloured based on their relationship to reconstructed pre-WGD chromosomes. Each set of bars represents chromosomes at different stages of acipenseriform evolution. The ancestral acipenseriform had 31 chromosomes. Through WGD, followed by chromosomal fusion, fission, and rearrangement events, the modern sturgeon genome now has 60 chromosomes. The plot shows 59 scaffolds based on the annotation from Du et al. (2020) with filtering steps applied to remove small scaffolds with few genes (see ‘Methods’). Some chromosomal segments experienced irreversible mixing and their origin could not be traced back to a single ancestral chromosome. This shared evolutionary history is represented by coloured stripes on sturgeon chromosomes.

**Figure S4:**
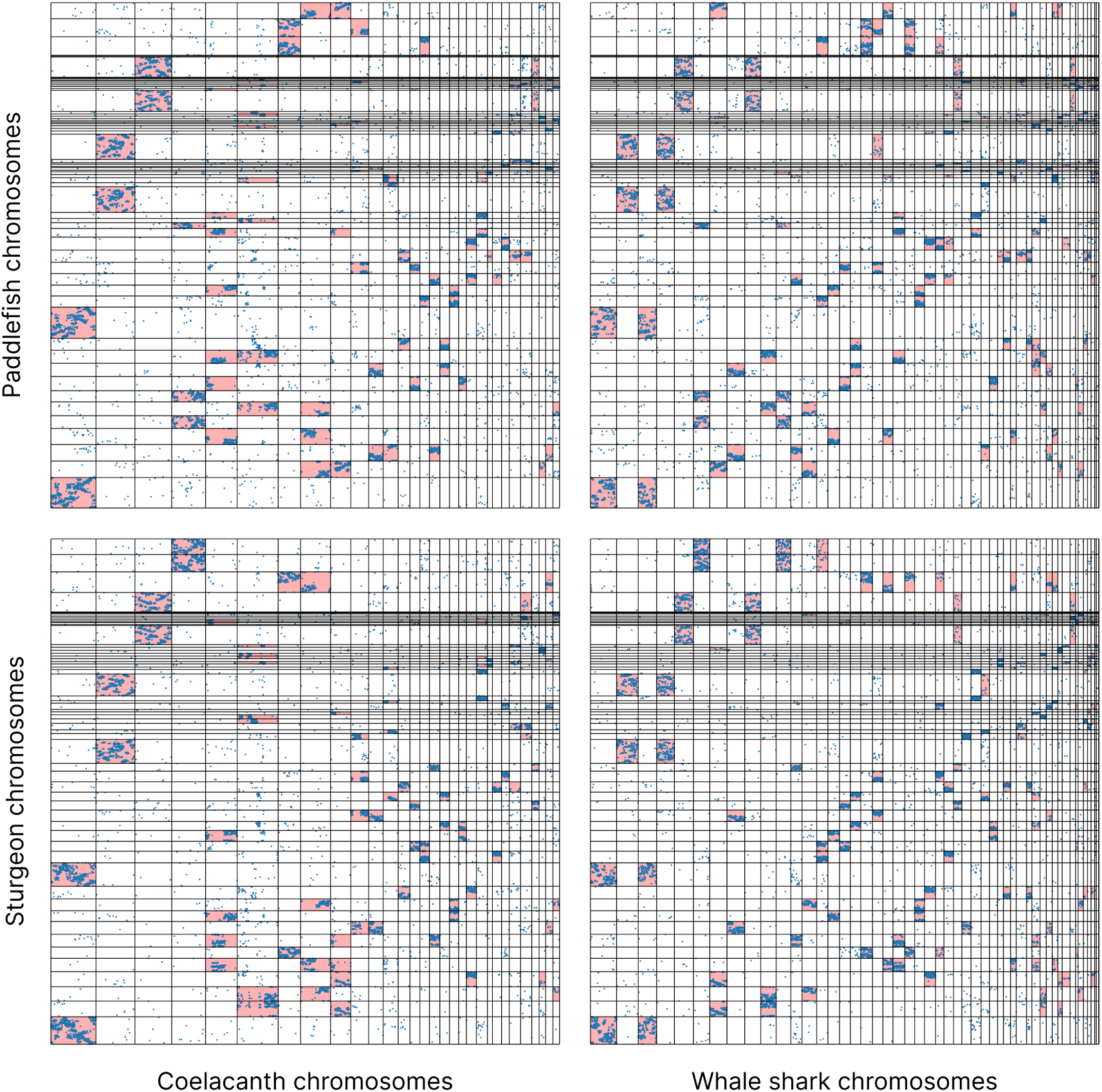
Macrosynteny conservation between acipenseriformes and outgroup species. Extant acipenseriform chromosomes were compared against coelacanth and whale shark chromosomes (x-axis). The acipenseriform chromosomes are shown along the y-axis, and their boundaries are indicated by horizontal black lines. Similarly, the coelacanth and whale shark chromosomes are delineated by vertical black lines. The locations of orthologous genes between acipenseriform chromosomes and each of the outgroup genomes were plotted and are indicated by blue dots. Chromosome pairs for which the enrichment of orthologs is significant are highlighted in red (see ‘Methods’).

**Figure S5:**
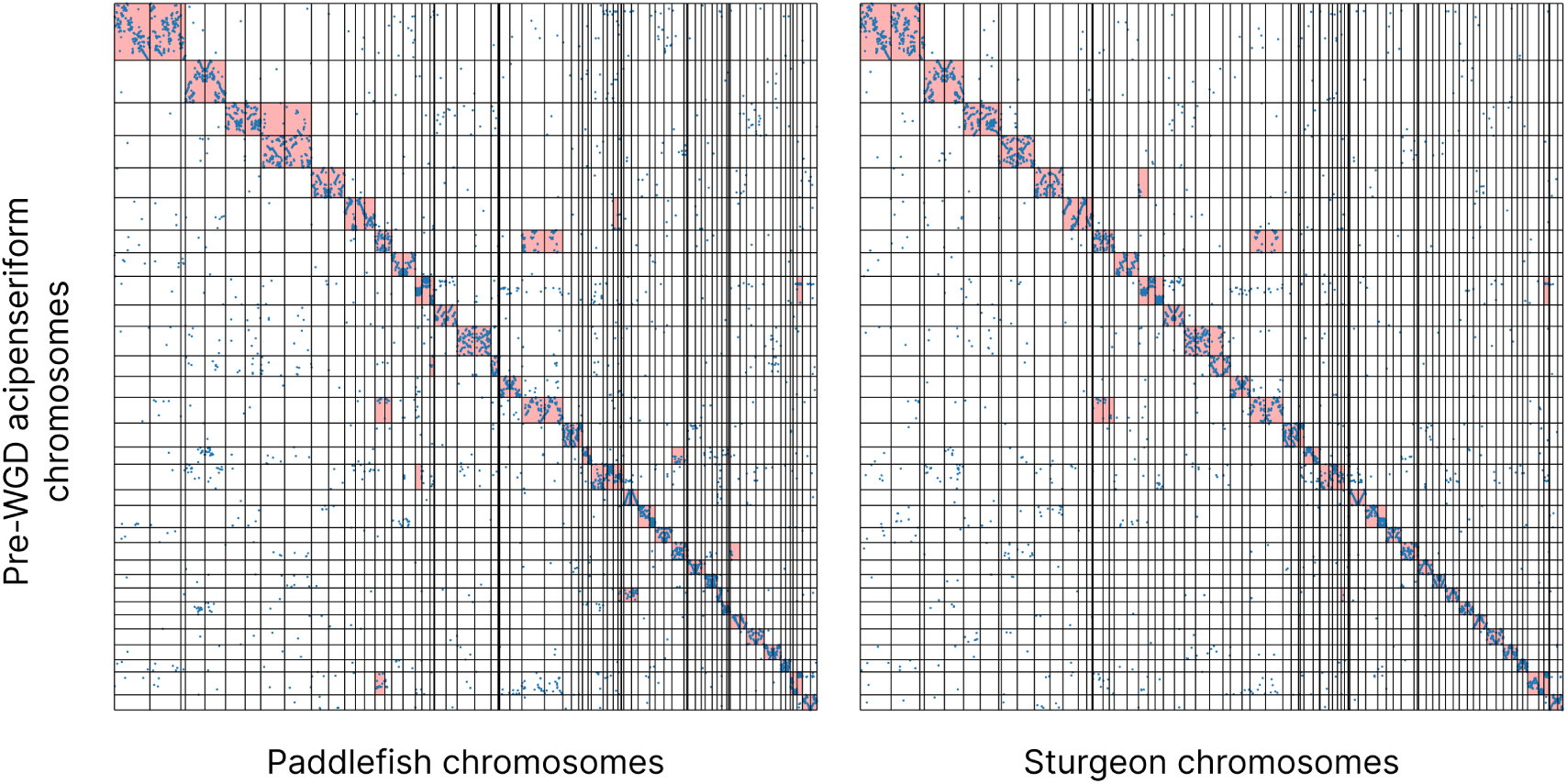
Distribution of orthologs between reconstructed pre-WGD chromosomes and extant acipenseriform genomes. Reconstructed pre-WGD acipenseriform chromosomes, as represented by spotted gar segments (y-axis), were compared against extant paddlefish and sturgeon chromosomes (x-axis). The reconstructed chromosomes are shown along the y-axis, and their boundaries are indicated by horizontal black lines. Similarly, the extant paddlefish and sturgeon chromosomes are delineated by vertical black lines. The locations of orthologous genes between reconstructed and extant chromosomes were plotted and are indicated by blue dots. Chromosome pairs for which the enrichment of orthologs is significant are highlighted in red (see ‘Methods’).

**Table S1:** Breakpoint consistency increases with additional segmentations. We segmented paddlefish and sturgeon against *K* genomes and compared these to segmentations with *K* + 1 genomes, recording the percentage of breakpoints that appeared in both segmentations. The percentage of breakpoints that remain consistent across segmentations increases with the number of genomes.

| **K** | **Paddlefish** | **Sturgeon** |
| --- | --- | --- |
| 1 | 55.84 | 58.68 |
| 2 | 61.43 | 60.71 |
| 3 | 77.59 | 78.75 |
| 4 | 78.64 | 75.41 |
| 5 | 90.02 | 90.87 |

**Table S2:** Significance of reconstruction for the acipenseriform ancestor. The significance of reconstructions containing *K* pre-WGD chromosomes. The log transformed probabilities calculated for paddlefish and sturgeon segments are shown (see Equation 3). The most significant number of ancestral acipenseriform chromosomes, *K* = 31, is highlighted in bold. The value of *K* was determined by comparing the sum of each row as both species did not converge on a single value.

| **K** | **Paddlefish** | **Sturgeon** |
| --- | --- | --- |
| 20 | -12713.27 | -17966.64 |
| 21 | -12865.44 | -18247.64 |
| 22 | -13238.17 | -18639.76 |
| 23 | -13369.77 | -18773.09 |
| 24 | -13472.74 | -18856.30 |
| 25 | -13610.82 | -19004.76 |
| 26 | -13716.58 | -19031.87 |
| 27 | -13838.08 | -19219.52 |
| 28 | -13923.78 | -19327.53 |
| 29 | -14018.82 | -19349.89 |
| 30 | -14140.01 | **-19482.19** |
| **31** | **-14346.32** | -19366.36 |
| 32 | -13039.60 | -19094.10 |
| 33 | -13038.11 | -19068.75 |
| 34 | -12872.77 | -19208.04 |
| 35 | -12282.65 | -18255.16 |
| 36 | -12419.24 | -18273.59 |
| 37 | -12514.32 | -18142.70 |
| 38 | -12587.13 | -18210.75 |
| 39 | -12658.44 | -18235.83 |
| 40 | -12656.81 | -17915.42 |

**Table S3:**
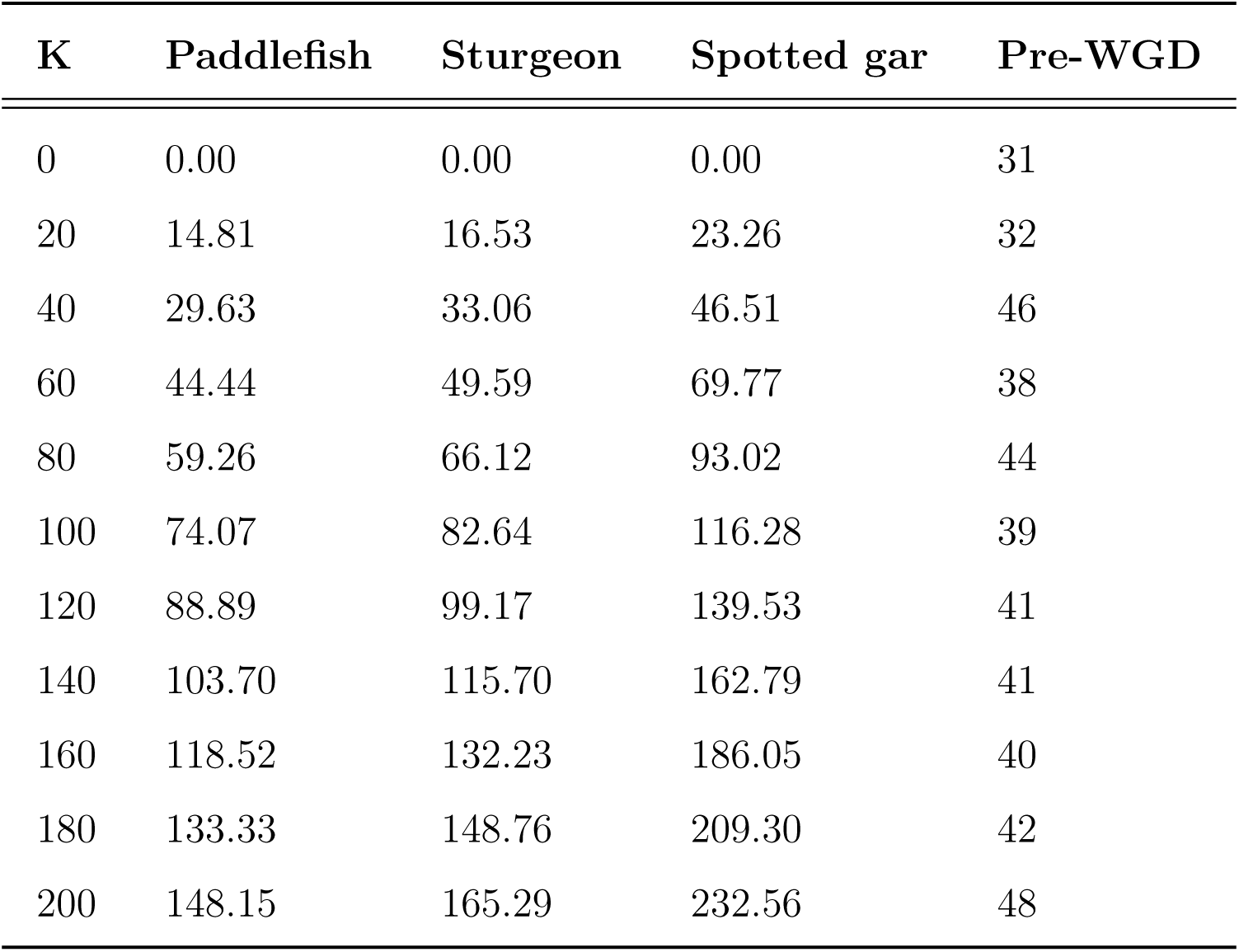
Reconstruction is consistent despite erroneous breakpoints. We introduced *K* random and erroneous genomic breakpoints into paddlefish, sturgeon, and spotted gar genomes, repeating the pre-WGD reconstruction for each value of *K*. The percentage of additional breakpoints in relation to our segmentations for paddlefish, sturgeon, and spotted gar are given for each species.

**Table S4:**
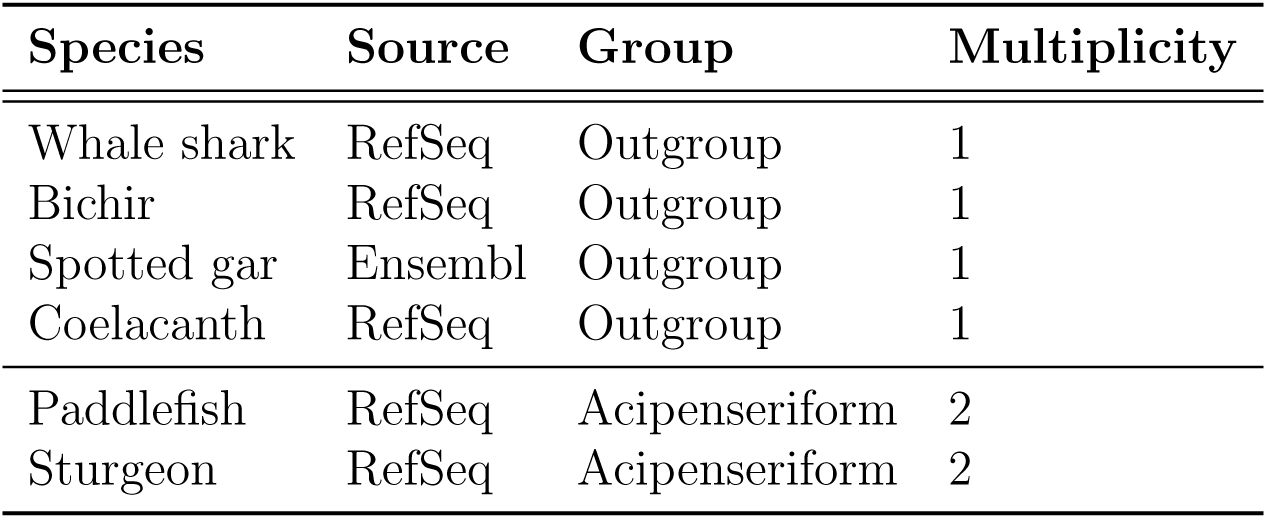
Species used in our reconstruction. Each species that was used in our analysis is listed below. These multiplicities are relative to the acipenseriform WGD.

| Species | Source | Group | Multiplicity |
| --- | --- | --- | --- |
| Whale shark | RefSeq | Outgroup | 1 |
| Bichir | RefSeq | Outgroup | 1 |
| Spotted gar | Ensembl | Outgroup | 1 |
| Coelacanth | RefSeq | Outgroup | 1 |
| Paddlefish | RefSeq | Acipenseriform | 2 |
| Sturgeon | RefSeq | Acipenseriform | 2 |

**Table S5:**
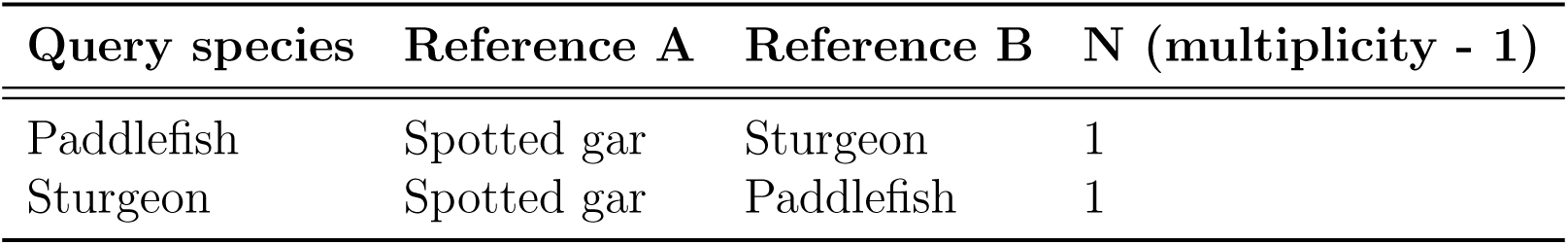
Genome trios used for paralogy inference. For each group, the query genome and two references are shown. Only paralogs that are dated between the divergence of reference A and reference B are included. This should allow us to discard lineage-specific duplications. The number of top hits (N) that we keep for each trio is displayed. This reflects the number of duplicates we expect given the whole genome duplication history of the query species.

| Query species | Reference A | Reference B | N (multiplicity - 1) |
| --- | --- | --- | --- |
| Paddlefish | Spotted gar | Sturgeon | 1 |
| Sturgeon | Spotted gar | Paddlefish | 1 |

**Table S6:** Breakpoint detection rate. For each cell in the table we generated 1000 sequences of different lengths where each element of the sequence was either a ‘1’ or ‘0’. These represent the presence or absence, respectively, of orthologs. We placed a breakpoint in the middle of each sequence such that the probability of observing a ‘1’ changes at this point. The difference in probability of observing a ‘1’ on either side of the breakpoint is given by the ‘Distance’ column. For example, a distance of ‘0.5’ means that the probability of observing a ‘1’ on the left of the breakpoint is 50% lower or higher compared to that on the right side of the breakpoint.

| Distance | Sequence length |  |  |  |
| --- | --- | --- | --- | --- |
|  | 10 | 20 | 40 | 80 |
| 1.00 | 1.00 | 1.00 | 1.00 | 1.00 |
| 0.90 | 0.91 | 0.91 | 0.92 | 0.95 |
| 0.80 | 0.85 | 0.86 | 0.86 | 0.91 |
| 0.70 | 0.73 | 0.82 | 0.86 | 0.88 |
| 0.60 | 0.65 | 0.78 | 0.84 | 0.91 |
| 0.50 | 0.55 | 0.76 | 0.85 | 0.88 |
| 0.40 | 0.42 | 0.69 | 0.81 | 0.86 |
| 0.30 | 0.30 | 0.57 | 0.73 | 0.84 |
| 0.20 | 0.18 | 0.42 | 0.66 | 0.78 |
| 0.10 | 0.09 | 0.22 | 0.42 | 0.66 |
| 0.00 | 0.00 | 0.00 | 0.00 | 0.00 |

**Table S7:** Breakpoint false-positive rate. For each cell in the table we generated 1000 sequences of different lengths where each element of the sequence was either a ‘1’ or ‘0’. These represent the presence or absence, respectively, of orthologs. The probability of observing a ‘1’ at any position of the sequence is given by the ‘Frequency’ column. Importantly, this does not change throughout the sequence and any breakpoints detected are erroneous.

| Frequency | Sequence length |  |  |  |
| --- | --- | --- | --- | --- |
|  | 10 | 20 | 40 | 80 |
| 1.00 | 0.00 | 0.00 | 0.00 | 0.00 |
| 0.90 | 0.01 | 0.03 | 0.03 | 0.02 |
| 0.80 | 0.03 | 0.05 | 0.03 | 0.03 |
| 0.70 | 0.07 | 0.07 | 0.05 | 0.05 |
| 0.60 | 0.07 | 0.06 | 0.04 | 0.03 |
| 0.50 | 0.07 | 0.07 | 0.02 | 0.04 |
| 0.40 | 0.07 | 0.06 | 0.04 | 0.03 |
| 0.30 | 0.07 | 0.05 | 0.04 | 0.05 |
| 0.20 | 0.03 | 0.06 | 0.05 | 0.04 |
| 0.10 | 0.01 | 0.02 | 0.04 | 0.02 |
| 0.00 | 0.00 | 0.00 | 0.00 | 0.00 |

## Notes

### Competing Interest Statement

The authors have declared no competing interest.

## References

C. T. Amemiya, J. Alföldi, A. P. Lee, S. Fan, H. Philippe, I. MacCallum, I. Braasch, T. Manousaki, I. Schneider, N. Rohner, C. Organ, D. Chalopin, J. J. Smith, M. Robinson, R. A. Dorrington, M. Gerdol, B. Aken, M. A. Biscotti, M. Barucca, D. Baurain, A. M. Berlin, G. L. Blatch, F. Buonocore, T. Burmester, M. S. Campbell, A. Canapa, J. P. Cannon, A. Christoffels, G. De Moro, A. L. Edkins, L. Fan, A. M. Fausto, N. Feiner, M. Forconi, J. Gamieldien, S. Gnerre, A. Gnirke, J. V. Goldstone, W. Haerty, M. E. Hahn, U. Hesse, S. Hoffmann, J. Johnson, S. I. Karchner, S. Kuraku, M. Lara, J. Z. Levin, G. W. Litman, E. Mauceli, T. Miyake, M. G. Mueller, D. R. Nelson, A. Nitsche, E. Olmo, T. Ota, A. Pallavicini, S. Panji, B. Picone, C. P. Ponting, S. J. Prohaska, D. Przybylski, N. R. Saha, V. Ravi, F. J. Ribeiro, T. Sauka-Spengler, G. Scapigliati, S. M. J. Searle, T. Sharpe, O. Simakov, P. F. Stadler, J. J. Stegeman, K. Sumiyama, D. Tabbaa, H. Tafer, J. Turner-Maier, P. van Heusden, S. White, L. Williams, M. Yandell, H. Brinkmann, J.-N. Volff, C. J. Tabin, N. Shubin, M. Schartl, D. B. Jaffe, J. H. Postlethwait, B. Venkatesh, F. Di Palma, E. S. Lander, A. Meyer, and K. Lindblad-Toh. The African coelacanth genome provides insights into tetrapod evolution. Nature, 496(7445):311–316, Apr. 2013. ISSN 1476-4687. doi: 10.1038/nature12027. URL https://www.nature.com/articles/nature12027.

J. Armstrong, G. Hickey, M. Diekhans, I. T. Fiddes, A. M. Novak, A. Deran, Q. Fang, D. Xie, S. Feng, J. Stiller, D. Genereux, J. Johnson, V. D. Marinescu, J. Alföldi, R. S. Harris, K. Lindblad-Toh, D. Haussler, E. Karlsson, E. D. Jarvis, G. Zhang, and B. Paten. Progressive Cactus is a multiple-genome aligner for the thousand-genome era. Nature, 587(7833):246–251, Nov. 2020. ISSN 1476-4687. doi: 10.1038/s41586-020-2871-y. URL https://www.nature.com/articles/s41586-020-2871-y.

I. Auger and C. Lawrence. Algorithms for the optimal identification of segment neighborhoods. Bulletin of Mathematical Biology, 51(1):39–54, 1989. ISSN 00928240. doi: 10.1016/S0092-8240(89)80047-3. URL http://link.springer.com/10.1016/S0092-8240(89)80047-3.

C. Bernard, Y. Nevers, N. B. R. Karampudi, K. J. Gilbert, C. Train, A. Warwick Vesztrocy, N. Glover, A. Altenhoff, and C. Dessimoz. EdgeHOG: a method for fine-grained ancestral gene order inference at large scale. Nature Ecology & Evolution, 9(10):1951–1961, Aug. 2025. ISSN 2397-334X. doi: 10.1038/s41559-025-02818-0. URL https://www.nature.com/articles/s41559-025-02818-0.

C. Berthelot, F. Brunet, D. Chalopin, A. Juanchich, M. Bernard, B. Nöel, P. Bento, C. Da Silva, K. Labadie, A. Alberti, J.-M. Aury, A. Louis, P. Dehais, P. Bardou, J. Montfort, C. Klopp, C. Cabau, C. Gaspin, G. H. Thorgaard, M. Boussaha, E. Quillet, R. Guyomard, D. Galiana, J. Bobe, J.-N. Volff, C. Genêt, P. Wincker, O. Jaillon, H. R. Crollius, and Y. Guiguen. The rainbow trout genome provides novel insights into evolution after whole-genome duplication in vertebrates. Nature Communications, 5 (1):3657, Apr. 2014. ISSN 2041-1723. doi: 10.1038/ncomms4657. URL https://www.nature.com/articles/ncomms4657.

X. Bi, K. Wang, L. Yang, H. Pan, H. Jiang, Q. Wei, M. Fang, H. Yu, C. Zhu, Y. Cai, Y. He, X. Gan, H. Zeng, D. Yu, Y. Zhu, H. Jiang, Q. Qiu, H. Yang, Y. E. Zhang, W. Wang, M. Zhu, S. He, and G. Zhang. Tracing the genetic footprints of vertebrate landing in non-teleost ray-finned fishes. Cell, 184(5):1377–1391.e14, Mar. 2021. ISSN 1097-4172. doi: 10.1016/j.cell.2021.01.046.

V. Birstein, R. Hanner, and R. Desalle. Phylogeny of the acipenseriformes: Cytogenetic and molecular approaches. Environmental Biology of Fishes, 48:127–155, 01 1997. doi: 10.1023/A:1007366100353.

I. Braasch, A. R. Gehrke, J. J. Smith, K. Kawasaki, T. Manousaki, J. Pasquier, A. Amores, T. Desvignes, P. Batzel, J. Catchen, A. M. Berlin, M. S. Campbell, D. Barrell, K. J. Martin, J. F. Mulley, V. Ravi, A. P. Lee, T. Nakamura, D. Chalopin, S. Fan, D. Wcisel, C. Canestro, J. Sydes, F. E. G. Beaudry, Y. Sun, J. Hertel, M. J. Beam, M. Fasold, M. Ishiyama, J. Johnson, S. Kehr, M. Lara, J. H. Letaw, G. W. Litman, R. T. Litman, M. Mikami, T. Ota, N. R. Saha, L. Williams, P. F. Stadler, H. Wang, J. S. Taylor, Q. Fontenot, A. Ferrara, S. M. J. Searle, B. Aken, M. Yandell, I. Schneider, J. A. Yoder, J.-N. Volff, A. Meyer, C. T. Amemiya, B. Venkatesh, P. W. H. Holland, Y. Guiguen, J. Bobe, N. H. Shubin, F. Di Palma, J. Alföldi, K. Lindblad-Toh, and J. H. Postlethwait. The spotted gar genome illuminates vertebrate evolution and facilitates human-teleost comparisons. Nature Genetics, 48(4):427–437, Apr. 2016. ISSN 1546-1718. doi: 10. 1038/ng.3526. URL https://www.nature.com/articles/ng.3526.

B. Buchfink, K. Reuter, and H.-G. Drost. Sensitive protein alignments at tree-of-life scale using DIAMOND. Nature Methods, 18(4):366–368, Apr. 2021. ISSN 1548-7105. doi: 10.1038/s41592-021-01101-x. URL https://www.nature.com/articles/s41592-021-01101-x.

D. Casey, Ą. Niezabitowski, M. Kumar Gundappa, H. Matz, A. Venugopalan, L. A. Hanson, H. Dooley, D. J. Macqueen, A. K. Redmond, and A. McLysaght. A duplicate resolved paddlefish genome provides insights into the mechanisms of rediploidisation and hox cluster evolution. Manuscript submitted for publication, submitted.

P. Cheng, Y. Huang, Y. Lv, H. Du, Z. Ruan, C. Li, H. Ye, H. Zhang, J. Wu, C. Wang, R. Ruan, Y. Li, C. Bian, X. You, C. Shi, K. Han, J. Xu, Q. Shi, and Q. Wei. The American Paddlefish Genome Provides Novel Insights into Chromosomal Evolution and Bone Mineralization in Early Vertebrates. Molecular Biology and Evolution, 38(4): 1595–1607, Apr. 2021. ISSN 1537-1719. doi: 10.1093/molbev/msaa326. URL https://academic.oup.com/mbe/article/38/4/1595/6040740.

J. W. Clark and P. C. Donoghue. Whole-Genome Duplication and Plant Macroevolution. Trends in Plant Science, 23(10):933–945, Oct. 2018. ISSN 13601385. doi: 10.1016/j.tplants.2018.07.006. URL https://linkinghub.elsevier.com/retrieve/ pii/S1360138518301596.

G. C. Conant and K. H. Wolfe. Turning a hobby into a job: how duplicated genes find new functions. Nature Reviews Genetics, 9(12):938–950, 2008.

P. Dehal and J. L. Boore. Two Rounds of Whole Genome Duplication in the Ancestral Vertebrate. PLOS Biology, 3(10):e314, Sept. 2005. ISSN 1545-7885. doi: 10.1371/ journal.pbio.0030314. URL https://journals.plos.org/plosbiology/article?id=10.1371/journal.pbio.0030314.

K. Du, M. Stöck, S. Kneitz, C. Klopp, J. M. Woltering, M. C. Adolfi, R. Feron, D. Prokopov, A. Makunin, I. Kichigin, C. Schmidt, P. Fischer, H. Kuhl, S. Wuertz, J. Gessner, W. Kloas, C. Cabau, C. Iampietro, H. Parrinello, C. Tomlinson, L. Journot, J. H. Postlethwait, I. Braasch, V. Trifonov, W. C. Warren, A. Meyer, Y. Guiguen, and M. Schartl. The sterlet sturgeon genome sequence and the mechanisms of segmental rediploidization. Nature Ecology & Evolution, 4(6):841–852, June 2020. ISSN 2397-334X. doi: 10.1038/s41559-020-1166-x. URL https://www.nature.com/articles/ s41559-020-1166-x.

K. Eilbeck, S. E. Lewis, C. J. Mungall, M. Yandell, L. Stein, R. Durbin, and M. Ashburner. The Sequence Ontology: a tool for the unification of genome annotations. Genome Biology, 6(5):R44, Apr. 2005. ISSN 1474-760X. doi: 10.1186/gb-2005-6-5-r44. URL 10.1186/gb-2005-6-5-r44.

D. M. Emms and S. Kelly. STRIDE: Species Tree Root Inference from Gene Duplication Events. Molecular Biology and Evolution, 34(12):3267–3278, Dec. 2017. ISSN 0737-4038, 1537-1719. doi: 10.1093/molbev/msx259. URL https://academic.oup.com/mbe/article/34/12/3267/4259048.

F. Fontana, L. Zane, A. Pepe, and L. Congiu. Polyploidy in acipenseriformes: Cytogenetic and molecular approaches. pages 385–403, 01 2007.

C. Fried, S. J. Prohaska, and P. F. Stadler. Independent Hox-cluster duplications in lampreys. Journal of Experimental Zoology Part B: Molecular and Developmental Evolution, 299B(1):18–25, Oct. 2003. ISSN 1552-5007, 1552-5015. doi: 10.1002/jez.b.37. URL https://onlinelibrary.wiley.com/doi/10.1002/jez.b.37.

R. F. Furlong, R. Younger, M. Kasahara, R. Reinhardt, M. Thorndyke, and P. W. H. Holland. A Degenerate ParaHox Gene Cluster in a Degenerate Vertebrate. Molecular Biology and Evolution, 24(12):2681–2686, June 2007. ISSN 0737-4038, 1537-1719. doi: 10.1093/molbev/msm194. URL https://academic.oup.com/mbe/article-lookup/ doi/10.1093/molbev/msm194.

W. R. Gilks. Introducing markov chain monte carlo. Markov chain Monte Carlo in practice, 1996.

M. Havelka, V. Kspar, M. Hulák, and M. Flaǰshans. Sturgeon genetics and cytogenetics: a review related to ploidy levels and interspecific hybridization. Folia Zoologica, 60(2): 93–103, June 2011. ISSN 0139-7893. doi: 10.25225/fozo.v60.i2.a3.2011. URL http://www.bioone.org/doi/10.25225/fozo.v60.i2.a3.2011.

Z. Huang, Z. Xu, H. Bai, Y. Huang, N. Kang, X. Ding, J. Liu, H. Luo, C. Yang, W. Chen, Q. Guo, L. Xue, X. Zhang, L. Xu, M. Chen, H. Fu, Y. Chen, Z. Yue, T. Fukagawa, S. Liu, G. Chang, and L. Xu. Evolutionary analysis of a complete chicken genome. Proceedings of the National Academy of Sciences, 120(8):e2216641120, Feb. 2023. ISSN 0027-8424, 1091-6490. doi: 10.1073/pnas.2216641120. URL 10.1073/pnas.2216641120.

J. G. Inoue, M. Miya, K. Lam, B.-H. Tay, J. A. Danks, J. Bell, T. I. Walker, and B. Venkatesh. Evolutionary Origin and Phylogeny of the Modern Holocephalans (Chondrichthyes: Chimaeriformes): A Mitogenomic Perspective. Molecular Biology and Evolution, 27(11):2576–2586, Nov. 2010. ISSN 0737-4038, 1537-1719. doi: 10.1093/molbev/msq147. URL https://academic.oup.com/mbe/article-lookup/ doi/10.1093/molbev/msq147.

Y. W. Kawaguchi, R. Matsumoto, and S. Kuraku. Improved genome assembly of whale shark, the world’s biggest fish: revealing intragenomic heterogeneity in molecular evolution. GigaScience, 15:giag014, Jan. 2026. ISSN 2047-217X. doi: 10.1093/gigascience/giag014. URL https://academic.oup.com/gigascience/article/doi/10.1093/gigascience/giag014/8466405.

M. Kellis, B. W. Birren, and E. S. Lander. Proof and evolutionary analysis of ancient genome duplication in the yeast Saccharomyces cerevisiae. Nature, 428(6983):617–624, Apr. 2004. ISSN 1476-4687. doi: 10.1038/nature02424. URL https://www.nature.com/articles/nature02424.

D. M. Larkin, G. Pape, R. Donthu, L. Auvil, M. Welge, and H. A. Lewin. Breakpoint regions and homologous synteny blocks in chromosomes have different evolutionary histories. Genome Research, 19(5):770–777, May 2009. ISSN 1088-9051, 1549-5469. doi: 10.1101/gr.086546.108. URL https://genome.cshlp.org/content/19/5/770.

T. D. Lewin, I. J.-Y. Liao, and Y.-J. Luo. Annelid Comparative Genomics and the Evolution of Massive Lineage-Specific Genome Rearrangement in Bilaterians. Molecular Biology and Evolution, 41(9):msae172, Sept. 2024. ISSN 0737-4038, 1537-1719. doi: 10.1093/molbev/msae172. URL https://academic.oup.com/mbe/article/doi/10. 1093/molbev/msae172/7733614.

T. D. Lewin, I. J.-Y. Liao, M.-E. Chen, J. D. Bishop, P. W. Holland, and Y.-J. Luo. Fusion, fission, and scrambling of the bilaterian genome in Bryozoa. Genome Research, 35(1):78–92, Jan. 2025. ISSN 1088-9051, 1549-5469. doi: 10.1101/gr.279636.124. URL http://genome.cshlp.org/lookup/doi/10.1101/gr.279636.124.

S. Lien, B. F. Koop, S. R. Sandve, J. R. Miller, M. P. Kent, T. Nome, T. R. Hvidsten, J. S. Leong, D. R. Minkley, A. Zimin, F. Grammes, H. Grove, A. Gjuvsland, B. Walenz, R. A. Hermansen, K. von Schalburg, E. B. Rondeau, A. Di Genova, J. K. A. Samy, J. Olav Vik, M. D. Vigeland, L. Caler, U. Grimholt, S. Jentoft, D. Inge Våge, P. de Jong, T. Moen, M. Baranski, Y. Palti, D. R. Smith, J. A. Yorke, A. J. Nederbragt, A. Tooming-Klunderud, K. S. Jakobsen, X. Jiang, D. Fan, Y. Hu, D. A. Liberles, R. Vidal, P. Iturra, S. J. M. Jones, I. Jonassen, A. Maass, S. W. Omholt, and W. S. Davidson. The Atlantic salmon genome provides insights into rediploidization. Nature, 533(7602):200–205, May 2016. ISSN 1476-4687. doi: 10.1038/nature17164. URL https://www.nature.com/articles/nature17164.

J. S. Liu and C. E. Lawrence. Bayesian inference on biopolymer models. Bioinformatics, 15 (1):38–52, Jan. 1999. ISSN 1367-4811, 1367-4803. doi: 10.1093/bioinformatics/15.1.38. URL https://academic.oup.com/bioinformatics/article/15/1/38/218372.

J. Lu, P. Huang, J. Sun, and J. Liu. DupScan: predicting and visualizing vertebrate genome duplication database. Nucleic Acids Research, 51(D1):D906–D912, Jan. 2023. ISSN 0305-1048, 1362-4962. doi: 10.1093/nar/gkac718. URL https://academic.oup.com/nar/article/51/D1/D906/6677325.

A. Ludwig, N. M. Belfiore, C. Pitra, V. Svirsky, and I. Jenneckens. Genome Duplication Events and Functional Reduction of Ploidy Levels in Sturgeon (Acipenser, Huso and Scaphirhynchus). Genetics, 158(3):1203–1215, July 2001. ISSN 1943-2631. doi: 10.1093/genetics/158.3.1203. URL https://academic.oup.com/genetics/article/158/3/1203/6049512.

D. J. Macqueen and I. A. Johnston. A well-constrained estimate for the timing of the salmonid whole genome duplication reveals major decoupling from species diversification. Proceedings of the Royal Society B: Biological Sciences, 281(1778): 20132881, Mar. 2014. ISSN 0962-8452, 1471-2954. doi: 10.1098/rspb.2013.2881. URL https://royalsocietypublishing.org/doi/10.1098/rspb.2013.2881.

T. Mandáková and M. A. Lysak. Post-polyploid diploidization and diversification through dysploid changes. Current opinion in plant biology, 42:55–65, 2018.

F. Marlétaz, N. Timoshevskaya, V. A. Timoshevskiy, E. Parey, O. Simakov, D. Gavriouchkina, M. Suzuki, K. Kubokawa, S. Brenner, J. J. Smith, and D. S. Rokhsar. The hagfish genome and the evolution of vertebrates. Nature, 627(8005):811–820, Mar. 2024. ISSN 1476-4687. doi: 10.1038/s41586-024-07070-3. URL https://www.nature.com/articles/s41586-024-07070-3.

A. McLysaght, K. Hokamp, and K. H. Wolfe. Extensive genomic duplication during early chordate evolution. Nature Genetics, 31(2):200–204, June 2002. ISSN 1061-4036, 1546-1718. doi: 10.1038/ng884. URL https://www.nature.com/articles/ng884z.

T. K. Mehta, V. Ravi, S. Yamasaki, A. P. Lee, M. M. Lian, B.-H. Tay, S. Tohari, S. Yanai, A. Tay, S. Brenner, and B. Venkatesh. Evidence for at least six Hox clusters in the Japanese lamprey ( Lethenteron japonicum ). Proceedings of the National Academy of Sciences, 110(40):16044–16049, Oct. 2013. ISSN 0027-8424, 1091-6490. doi: 10.1073/pnas.1315760110. URL https://pnas.org/doi/full/10.1073/pnas.1315760110.

M. Muffato, A. Louis, N. T. T. Nguyen, J. Lucas, C. Berthelot, and H. Roest Crollius. Reconstruction of hundreds of reference ancestral genomes across the eukaryotic kingdom. Nature Ecology & Evolution, 7(3):355–366, Jan. 2023. ISSN 2397-334X. doi: 10.1038/s41559-022-01956-z. URL https://www.nature.com/articles/s41559-022-01956-z.

F. Mölder, K. P. Jablonski, B. Letcher, M. B. Hall, C. H. Tomkins-Tinch, V. Sochat, J. Forster, S. Lee, S. O. Twardziok, A. Kanitz, A. Wilm, M. Holtgrewe, S. Rahmann, S. Nahnsen, and J. Köster. Sustainable data analysis with Snakemake, Jan. 2021. URL https://f1000research.com/articles/10-33.

Y. Nakatani and A. McLysaght. Genomes as documents of evolutionary history: a probabilistic macrosynteny model for the reconstruction of ancestral genomes. Bioinformatics, 33(14):i369–i378, July 2017. ISSN 1367-4803, 1460-2059. doi: 10.1093/bioinformatics/btx259. URL https://academic.oup.com/bioinformatics/article/33/14/i369/3953974.

Y. Nakatani, H. Takeda, Y. Kohara, and S. Morishita. Reconstruction of the vertebrate ancestral genome reveals dynamic genome reorganization in early vertebrates. Genome Research, 17(9):1254–1265, Sept. 2007. ISSN 1088-9051. doi: 10.1101/gr.6316407. URL

http://genome.cshlp.org/lookup/doi/10.1101/gr.6316407.

Y. Nakatani, P. Shingate, V. Ravi, N. E. Pillai, A. Prasad, A. McLysaght, and B. Venkatesh. Reconstruction of proto-vertebrate, proto-cyclostome and proto-gnathostome genomes provides new insights into early vertebrate evolution. Nature Communications, 12(1):4489, July 2021. ISSN 2041-1723. doi: 10.1038/s41467-021-24573-z. URL https://www.nature.com/articles/s41467-021-24573-z.

L. Niezabitowski, R. Long, K. A. Redmond, and A. McLysaght. A comprehensive and accessible database of jawed-vertebrate ohnologs. Science Advances, forthcoming. Accepted for publication.

S. Ohno. *Evolution by Gene Duplication*. Springer Berlin Heidelberg, Berlin, Heidelberg, 1970. ISBN 9783642866616 9783642866593. doi: 10.1007/978-3-642-86659-3. URL http://link.springer.com/10.1007/978-3-642-86659-3.

E. Parey, A. Louis, J. Montfort, Y. Guiguen, H. R. Crollius, and C. Berthelot. An atlas of fish genome evolution reveals delayed rediploidization following the teleost whole-genome duplication. preprint, Genomics, Jan. 2022. URL http://biorxiv.org/lookup/doi/10.1101/2022.01.13.476171.

W. R. Pearson and D. J. Lipman. Improved tools for biological sequence comparison. Proceedings of the National Academy of Sciences, 85(8):2444–2448, Apr. 1988. ISSN 0027-8424, 1091-6490. doi: 10.1073/pnas.85.8.2444. URL https://pnas.org/doi/full/10.1073/pnas.85.8.2444.

S. Proost, J. Fostier, D. De Witte, B. Dhoedt, P. Demeester, Y. Van De Peer, and K. Vandepoele. i-ADHoRe 3.0—fast and sensitive detection of genomic homology in extremely large data sets. Nucleic Acids Research, 40(2):e11–e11, Jan. 2012. ISSN 1362-4962, 0305-1048. doi: 10.1093/nar/gkr955. URL https://academic.oup.com/nar/article/40/2/e11/2409742.

J. Rajkov, Z. Shao, and P. Berrebi. Evolution of Polyploidy and Functional Diploidization in Sturgeons: Microsatellite Analysis in 10 Sturgeon Species. Journal of Heredity, 105(4):521–531, July 2014. ISSN 0022-1503, 1465-7333. doi: 10.1093/jhered/esu027. URL https://academic.oup.com/jhered/article-lookup/doi/10.1093/jhered/esu027.

A. K. Redmond, D. Casey, M. K. Gundappa, D. J. Macqueen, and A. McLysaght. In-dependent rediploidization masks shared whole genome duplication in the sturgeon-paddlefish ancestor. Nature Communications, 14(1):2879, May 2023. ISSN 2041-1723. doi: 10.1038/s41467-023-38714-z. URL https://www.nature.com/articles/s41467-023-38714-z.

F. M. Robertson, M. K. Gundappa, F. Grammes, T. R. Hvidsten, A. K. Redmond, S. Lien, S. A. M. Martin, P. W. H. Holland, S. R. Sandve, and D. J. Macqueen. Lineage-specific rediploidization is a mechanism to explain time-lags between genome duplication and evolutionary diversification. Genome Biology, 18(1):111, June 2017. ISSN 1474-760X. doi: 10.1186/s13059-017-1241-z. URL 10.1186/s13059-017-1241-z.

C. Sacerdot, A. Louis, C. Bon, C. Berthelot, and H. Roest Crollius. Chromosome evolution at the origin of the ancestral vertebrate genome. Genome Biology, 19(1):166, Oct. 2018. ISSN 1474-760X. doi: 10.1186/s13059-018-1559-1. URL 10.1186/s13059-018-1559-1.

D. T. Schultz, S. H. D. Haddock, J. V. Bredeson, R. E. Green, O. Simakov, and D. S. Rokhsar. Ancient gene linkages support ctenophores as sister to other animals. Nature, 618(7963):110–117, June 2023. ISSN 0028-0836, 1476-4687. doi: 10.1038/s41586-023-05936-6. URL https://www.nature.com/articles/s41586-023-05936-6.

O. Simakov, F. Marlétaz, J.-X. Yue, B. O’Connell, J. Jenkins, A. Brandt, R. Calef, C.-H. Tung, T.-K. Huang, J. Schmutz, N. Satoh, J.-K. Yu, N. H. Putnam, R. E. Green, and D. S. Rokhsar. Deeply conserved synteny resolves early events in vertebrate evolution. Nature Ecology & Evolution, 4(6):820–830, June 2020. ISSN 2397-334X. doi: 10.1038/ s41559-020-1156-z. URL https://www.nature.com/articles/s41559-020-1156-z.

O. Simakov, J. Bredeson, K. Berkoff, F. Marletaz, T. Mitros, D. T. Schultz, B. L. O’Connell, P. Dear, D. E. Martinez, R. E. Steele, R. E. Green, C. N. David, and D. S. Rokhsar. Deeply conserved synteny and the evolution of metazoan chromosomes. Science Advances, 8(5):eabi5884, Feb. 2022. ISSN 2375-2548. doi: 10.1126/sciadv.abi5884. URL https://www.science.org/doi/10.1126/sciadv.abi5884.

P. P. Singh and H. Isambert. OHNOLOGS v2: a comprehensive resource for the genes retained from whole genome duplication in vertebrates. Nucleic Acids Research, page gkz909, Oct. 2019. ISSN 0305-1048, 1362-4962. doi: 10.1093/nar/gkz909. URL https://academic.oup.com/nar/advance-article/doi/10.1093/nar/gkz909/5587630.

P. P. Singh, J. Arora, and H. Isambert. Identification of Ohnolog Genes Originating from Whole Genome Duplication in Early Vertebrates, Based on Synteny Comparison across Multiple Genomes. PLOS Computational Biology, 11(7):e1004394, July 2015. ISSN 1553-7358. doi: 10.1371/journal.pcbi.1004394. URL 10.1371/journal.pcbi.1004394.

J. J. Smith, S. Kuraku, C. Holt, T. Sauka-Spengler, N. Jiang, M. S. Campbell, M. D. Yan-dell, T. Manousaki, A. Meyer, O. E. Bloom, J. R. Morgan, J. D. Buxbaum, R. Sachidanandam, C. Sims, A. S. Garruss, M. Cook, R. Krumlauf, L. M. Wiedemann, S. A. Sower, W. A. Decatur, J. A. Hall, C. T. Amemiya, N. R. Saha, K. M. Buckley, J. P. Rast, S. Das, M. Hirano, N. McCurley, P. Guo, N. Rohner, C. J. Tabin, P. Piccinelli, G. Elgar, M. Ruffier, B. L. Aken, S. M. J. Searle, M. Muffato, M. Pignatelli, J. Herrero, M. Jones, C. T. Brown, Y.-W. Chung-Davidson, K. G. Nanlohy, S. V. Libants, C.-Y. Yeh, D. W. McCauley, J. A. Langeland, Z. Pancer, B. Fritzsch, P. J. de Jong, B. Zhu, L. L. Fulton, B. Theising, P. Flicek, M. E. Bronner, W. C. Warren, S. W. Clifton, R. K. Wilson, and W. Li. Sequencing of the sea lamprey (Petromyzon marinus) genome provides insights into vertebrate evolution. Nature Genetics, 45(4):415–421, Apr. 2013. ISSN 1546-1718. doi: 10.1038/ng.2568. URL https://www.nature.com/articles/ng.2568.

J. J. Smith, N. Timoshevskaya, C. Ye, C. Holt, M. C. Keinath, H. J. Parker, M. E. Cook, J. E. Hess, S. R. Narum, F. Lamanna, H. Kaessmann, V. A. Timoshevskiy, C. K. M. Waterbury, C. Saraceno, L. M. Wiedemann, S. M. C. Robb, C. Baker, E. E. Eichler, D. Hockman, T. Sauka-Spengler, M. Yandell, R. Krumlauf, G. Elgar, and C. T. Amemiya. The sea lamprey germline genome provides insights into programmed genome rearrangement and vertebrate evolution. Nature Genetics, 50(2):270–277, Feb. 2018. ISSN 1546-1718. doi: 10.1038/s41588-017-0036-1. URL https://www.nature.com/articles/s41588-017-0036-1.

R. Symonová, M. Havelka, C. T. Amemiya, W. M. Howell, T. Kŕınková, M. Flaǰshans, D. Gela, and P. Ráb. Molecular cytogenetic differentiation of paralogs of Hox paralogs in duplicated and re-diploidized genome of the North American paddlefish (Polyodon spathula). BMC Genetics, 18(1):19, Mar. 2017. ISSN 1471-2156. doi: 10.1186/s12863-017-0484-8. URL 10.1186/s12863-017-0484-8.

The Arabidopsis Genome Initiative. Analysis of the genome sequence of the flowering plant Arabidopsis thaliana. *Nature*, 408(6814):796–815, Dec. 2000. ISSN 0028-0836, 1476-4687. doi: 10.1038/35048692. URL https://www.nature.com/articles/35048692.

V. A. Trifonov, S. S. Romanenko, V. R. Beklemisheva, L. S. Biltueva, A. I. Makunin, N. A. Lemskaya, A. I. Kulemzina, R. Stanyon, and A. S. Graphodatsky. Evolutionary plasticity of acipenseriform genomes. Chromosoma, 125(4):661–668, Sept. 2016. ISSN 0009-5915, 1432-0886. doi: 10.1007/s00412-016-0609-2. URL http://link.springer.com/10.1007/s00412-016-0609-2.

Y. Van de Peer, S. Maere, and A. Meyer. The evolutionary significance of ancient genome duplications. Nature Reviews Genetics, 10(10):725–732, Oct. 2009. ISSN 1471-0064. doi: 10.1038/nrg2600. URL https://www.nature.com/articles/nrg2600.

Y. Van de Peer, E. Mizrachi, and K. Marchal. The evolutionary significance of polyploidy. Nature Reviews Genetics, 18(7):411–424, July 2017. ISSN 1471-0064. doi: 10.1038/nrg.2017.26. URL https://www.nature.com/articles/nrg.2017.26.

C. Vargas-Chávez, L. Benıtez-Álvarez, G. I. Martınez-Redondo, L. Álvarez Gonźalez, J. Salces-Ortiz, K. Eleftheriadi, N. Escudero, N. Guiglielmoni, J.-F. Flot, M. Novo, A. Ruiz-Herrera, A. McLysaght, and R. Ferńandez. An episodic burst of massive genomic rearrangements and the origin of non-marine annelids. Nature Ecology & Evolution, 9(7):1263–1279, July 2025. ISSN 2397-334X. doi: 10.1038/s41559-025-02728-1. URL https://www.nature.com/articles/s41559-025-02728-1.

B. Venkatesh, A. P. Lee, V. Ravi, A. K. Maurya, M. M. Lian, J. B. Swann, Y. Ohta, M. F. Flajnik, Y. Sutoh, M. Kasahara, S. Hoon, V. Gangu, S. W. Roy, M. Irimia, V. Korzh, I. Kondrychyn, Z. W. Lim, B.-H. Tay, S. Tohari, K. W. Kong, S. Ho, B. Lorente-Galdos, J. Quilez, T. Marques-Bonet, B. J. Raney, P. W. Ingham, A. Tay, L. W. Hillier, P. Minx, T. Boehm, R. K. Wilson, S. Brenner, and W. C. Warren. Elephant shark genome provides unique insights into gnathostome evolution. Nature, 505(7482): 174–179, Jan. 2014. ISSN 1476-4687. doi: 10.1038/nature12826. URL https://www.nature.com/articles/nature12826.

J.-N. Volff. Genome evolution and biodiversity in teleost fish. Heredity, 94(3):280–294, Mar. 2005. ISSN 1365-2540. doi: 10.1038/sj.hdy.6800635. URL https://www.nature. com/articles/6800635.

Y. Wang, H. Tang, J. D. DeBarry, X. Tan, J. Li, X. Wang, T.-h. Lee, H. Jin, B. Marler, H. Guo, J. C. Kissinger, and A. H. Paterson. MCScanX: a toolkit for detection and evolutionary analysis of gene synteny and collinearity. Nucleic Acids Research, 40(7): e49–e49, Apr. 2012. ISSN 0305-1048, 1362-4962. doi: 10.1093/nar/gkr1293. URL https://academic.oup.com/nar/article-lookup/doi/10.1093/nar/gkr1293.

Y. Wang, H. Tang, X. Wang, Y. Sun, P. V. Joseph, and A. H. Paterson. Detection of colinear blocks and synteny and evolutionary analyses based on utilization of MCScanX. Nature Protocols, 19(7):2206–2229, July 2024. ISSN 1750-2799. doi: 10.1038/s41596-024-00968-2. URL https://www.nature.com/articles/s41596-024-00968-2.

K. H. Wolfe. Yesterday’s polyploids and the mystery of diploidization. Nature Reviews Genetics, 2(5):333–341, May 2001. ISSN 1471-0056, 1471-0064. doi: 10.1038/35072009. URL https://www.nature.com/articles/35072009.

K. H. Wolfe and D. C. Shields. Molecular evidence for an ancient duplication of the entire yeast genome. Nature, 387(6634):708–713, June 1997. ISSN 1476-4687. doi: 10.1038/42711. URL https://www.nature.com/articles/42711.

C. Xie, Z. Ma, C. Zhou, K. Ma, H. Wang, J. Wu, Y. Zhou, Y. Lu, D. Ji, X. Gu, H. Gao, J. Li, S. Fu, W. Li, Z. Han, S. Xiao, F. Liu, B. Zeng, S. Chen, J. Niu, T. Zhang, J. Shen, C. Liu, J. Luo, D. J. Macqueen, A. Meyer, H. Liu, and L. Xu. Chromosomal fusions trigger rediploidization of autopolyploid genomes. Nature, Apr. 2026. ISSN 1476-4687. doi: 10.1038/s41586-026-10439-1. URL 10.1038/s41586-026-10439-1.

A. D. Yates, P. Achuthan, W. Akanni, J. Allen, J. Allen, J. Alvarez-Jarreta, M. R. Amode, I. M. Armean, A. G. Azov, R. Bennett, J. Bhai, K. Billis, S. Boddu, J. C. Marugán, C. Cummins, C. Davidson, K. Dodiya, R. Fatima, A. Gall, C. G. Giron, L. Gil, T. Grego, L. Haggerty, E. Haskell, T. Hourlier, O. G. Izuogu, S. H. Janacek, T. Juettemann, M. Kay, I. Lavidas, T. Le, D. Lemos, J. G. Martinez, T. Maurel, M. McDowall, A. McMahon, S. Mohanan, B. Moore, M. Nuhn, D. N. Oheh, A. Parker, A. Parton, M. Patricio, M. P. Sakthivel, A. I. Abdul Salam, B. M. Schmitt, H. Schuilenburg, D. Sheppard, M. Sycheva, M. Szuba, K. Taylor, A. Thormann, G. Threadgold, A. Vullo, B. Walts, A. Winterbottom, A. Zadissa, M. Chakiachvili, B. Flint, A. Frankish, S. E. Hunt, G. IIsley, M. Kostadima, N. Langridge, J. E. Loveland, F. J. Martin, J. Morales, J. M. Mudge, M. Muffato, E. Perry, M. Ruffier, S. J. Trevanion, F. Cunningham, K. L. Howe, D. R. Zerbino, and P. Flicek. Ensembl 2020. *Nucleic Acids Research*, page gkz966, Nov. 2019. ISSN 0305-1048, 1362-4962. doi: 10.1093/nar/gkz966. URL https://academic.oup.com/nar/advance-article/doi/10.1093/nar/gkz966/5613682.

D. Yu, Y. Ren, M. Uesaka, A. J. S. Beavan, M. Muffato, J. Shen, Y. Li, I. Sato, W. Wan, J. W. Clark, J. N. Keating, E. M. Carlisle, R. P. Dearden, S. Giles, E. Randle, R. S. Sansom, R. Feuda, J. F. Fleming, F. Sugahara, C. Cummins, M. Patricio, W. Akanni, S. D’Aniello, C. Bertolucci, N. Irie, C. Alev, G. Sheng, A. de Mendoza, I. Maeso, M. Irimia, B. Fromm, K. J. Peterson, S. Das, M. Hirano, J. P. Rast, M. D. Cooper, J. Paps, D. Pisani, S. Kuratani, F. J. Martin, W. Wang, P. C. J. Donoghue, Y. E. Zhang, and J. Pascual-Anaya. Hagfish genome elucidates vertebrate whole-genome duplication events and their evolutionary consequences. Nature Ecology & Evolution, 8(3):519–535, Mar. 2024. ISSN 2397-334X. doi: 10.1038/s41559-023-02299-z. URL https://www.nature.com/articles/s41559-023-02299-z.

## Supplementary References

K. Du, M. Stöck, S. Kneitz, C. Klopp, J. M. Woltering, M. C. Adolfi, R. Feron, D. Prokopov, A. Makunin, I. Kichigin, C. Schmidt, P. Fischer, H. Kuhl, S. Wuertz, J. Gessner, W. Kloas, C. Cabau, C. Iampietro, H. Parrinello, C. Tomlinson, L. Journot, J. H. Postlethwait, I. Braasch, V. Trifonov, W. C. Warren, A. Meyer, Y. Guiguen, and M. Schartl. The sterlet sturgeon genome sequence and the mechanisms of segmental rediploidization. Nature Ecology & Evolution, 4(6):841–852, June 2020. ISSN 2397-334X. doi: 10.1038/s41559-020-1166-x. URL https://www.nature.com/articles/s41559-020-1166-x.

M. K. Gundappa, T.-H. To, L. Grønvold, S. A. M. Martin, S. Lien, J. Geist, D. Hazlerigg, S. R. Sandve, and D. J. Macqueen. Genome-Wide Reconstruction of Rediploidization Following Autopolyploidization across One Hundred Million Years of Salmonid Evolution. Molecular Biology and Evolution, 39(1):msab310, Jan. 2022. ISSN 0737-4038, 1537-1719. doi: 10.1093/molbev/msab310. URL https://academic.oup.com/mbe/article/doi/10.1093/molbev/msab310/6413642.

A. K. Redmond, D. Casey, M. K. Gundappa, D. J. Macqueen, and A. McLysaght. Independent rediploidization masks shared whole genome duplication in the sturgeon-paddlefish ancestor. Nature Communications, 14(1):2879, May 2023. ISSN 2041-1723. doi: 10.1038/s41467-023-38714-z. URL https://www.nature.com/articles/s41467-023-38714-z.

